# A common haplotype lowers PU.1 expression in myeloid cells and delays onset of Alzheimer’s disease

**DOI:** 10.1101/110957

**Authors:** Kuan-lin Huang, Edoardo Marcora, Anna A Pimenova, Antonio F Di Narzo, Manav Kapoor, Sheng Chih Jin, Oscar Harari, Sarah Bertelsen, Benjamin P Fairfax, Jake Czajkowski, Vincent Chouraki, Benjamin Grenier-Boley, Céline Bellenguez, Yuetiva Deming, Andrew McKenzie, Towfique Raj, Alan E Renton, John Budde, Albert Smith, Annette Fitzpatrick, Joshua C Bis, Anita DeStefano, Hieab HH Adams, M Arfan Ikram, Sven van der Lee, Jorge L. Del-Aguila, Maria Victoria Fernandez, Laura Ibañez, The International Genomics of Alzheimer’s Project, The Alzheimer’s Disease Neuroimaging Initiative, Rebecca Sims, Valentina Escott-Price, Richard Mayeux, Jonathan L Haines, Lindsay A Farrer, Margaret A. Pericak-Vance, Jean Charles Lambert, Cornelia van Duijn, Lenore Launer, Sudha Seshadri, Julie Williams, Philippe Amouyel, Gerard D Schellenberg, Bin Zhang, Ingrid Borecki, John S K Kauwe, Carlos Cruchaga, Ke Hao, Alison M Goate

## Abstract

A genome-wide survival analysis of 14,406 Alzheimer’s disease (AD) cases and 25,849 controls identified eight previously reported AD risk loci and fourteen novel loci associated with age at onset. LD score regression of 220 cell types implicated regulation of myeloid gene expression in AD risk. In particular, the minor allele of rs1057233 (G), within the previously reported *CELF1* AD risk locus, showed association with delayed AD onset and lower expression of *SPI1* in monocytes and macrophages. *SPI1* encodes PU.1, a transcription factor critical for myeloid cell development and function. AD heritability is enriched within the PU.1 cistrome, implicating a myeloid PU.1 target gene network in AD. Finally, experimentally altered PU.1 levels affect the expression of mouse orthologs of many AD risk genes and the phagocytic activity of mouse microglial cells. Our results suggest that lower *SPI1* expression reduces AD risk by regulating myeloid gene expression and cell function.

---

AD is the most prevalent form of dementia. While genome-wide association studies (GWAS) have identified more than twenty AD risk loci^1–5^, the associated disease genes and mechanisms remain largely unclear. To better understand these genetic associations, AD-related phenotypes can be leveraged. For example, few studies^6,7^ have investigated the genetic basis of age at onset of AD (AAO). To date, *APOE* remains the only locus repeatedly associated with AAO^8,9^, but *PICALM* and *BIN1* have also been reported to affect AAO^6,10,11^. Further, we have previously used CSF biomarkers to demonstrate that *APOE* genotype is strongly associated with these disease-relevant endophenotypes^12,13^.

Identifying causal genes and mechanisms underlying disease-associated loci requires integrative analyses of expression and epigenetic datasets in disease-relevant cell types^14^. Recent genetic and molecular evidence has highlighted the role of myeloid cells in AD pathogenesis. At the genetic level, GWAS and sequencing studies have found associations between AD and genes expressed in myeloid cells, including *TREM2*, *ABCA7*, and *CD33*^1,2,5,15–17^. At the epigenetic level, genes expressed in myeloid cells display abnormal patterns of chromatin modification in AD mouse models and human samples^18–20^. Further, AD-risk alleles are polarized for *cis*-expression quantitative trait locus (*cis*-eQTL) effects in monocytes^21^. Herein, we show that AD heritability is enriched in functional annotations for cells of the myeloid and B-lymphoid lineage, suggesting that integrative analyses of AD loci with myeloid-specific expression and epigenetic datasets will uncover novel AD genes and mechanisms related to the function of these cell types.

In this study, we conducted a large-scale genome-wide survival analysis and subsequent endophenotype association analysis to uncover loci associated with AAO-defined survival (AAOS) in AD cases and non-demented elderly controls. We discovered an AAOS- and CSF Aβ_42_-associated SNP, rs1057233, in the previously reported *CELF1* AD risk locus. *Cis*-eQTL analyses revealed a highly significant association of the protective rs1057233^G^ allele with reduced *SPI1* expression in human myeloid cells. *SPI1* encodes PU.1, a transcription factor critical for myeloid and B-lymphoid cell development and function, that binds to the *cis*-regulatory elements of several AD-associated genes in these cells. Moreover, we show that AD heritability is enriched in PU.1 ChIP-Seq binding sites in human myeloid cells across the genome, implicating a myeloid PU.1 target gene network in the etiology of AD. To validate these bioinformatic analyses, we show that experimentally altered PU.1 levels are correlated with phagocytic activity of mouse microglial cells and the expression of multiple genes involved in diverse biological processes of myeloid cells. This evidence collectively shows that lower *SPI1* expression may reduce AD risk by modulating myeloid cell gene expression and function.

## Results

### Genome-wide survival analysis

For the genome-wide survival analysis, we used 14,406 AD case and 25,849 control samples from the IGAP consortium (Table 1a). 8,253,925 SNPs passed quality control and were included for meta-analysis across all cohorts (**Supplementary Table 1**), which showed little evidence of genomic inflation (λ=1.026). Four loci showed genome-wide significant associations (P <5×10^−8^) with AAOS: *BIN1* (P=7.6×10^−13^), *MS4A* (P=5.1×10^−11^), *PICALM* (P=4.3×10^−14^), and *APOE* (P=1.2×10^−67^) (Supplementary Fig. 1). While SNPs within *BIN1*^6^, *PICALM*^6,10^, and *APOE*^6,8–10,22^ loci have previously been shown to be associated with AAO, this is the first time that the *MS4A* locus is reported to be associated with an AAO-related phenotype. The minor allele of rs7930318 near *MS4A4A* is associated with delayed AAO. Four other AD risk loci previously reported in the IGAP GWAS^1^ showed associations that reached suggestive significance (P <1.0×10^−5^): *CR1* (P=1.2×10^−6^), *SPI1/CELF1* (P=5.4×10^−6^), *SORL1* (P=1.8×10^−7^), and *FERMT2* (P=1.0×10^−5^). The direction of effects were concordant with the previous IGAP GWAS logistic regression analysis for AD risk^1^ at all suggestive loci: AD risk-increasing alleles were all associated with a hazard ratio above 1 and earlier AAO, whereas AD risk-decreasing alleles were all associated with a hazard ratio below 1 and later AAO (Table 1b, **Supplementary Table 2**). We also identified 14 novel loci that reached suggestive significance in the survival analysis, 3 of which (rs116341973, rs1625716, and rs11074412) were nominally associated with AD risk (Bonferroni-corrected threshold: P=0.05/22=2.27×10^−3^) in the IGAP GWAS (Table 1b, Supplementary Fig. 2, 3).

**Table 1.**
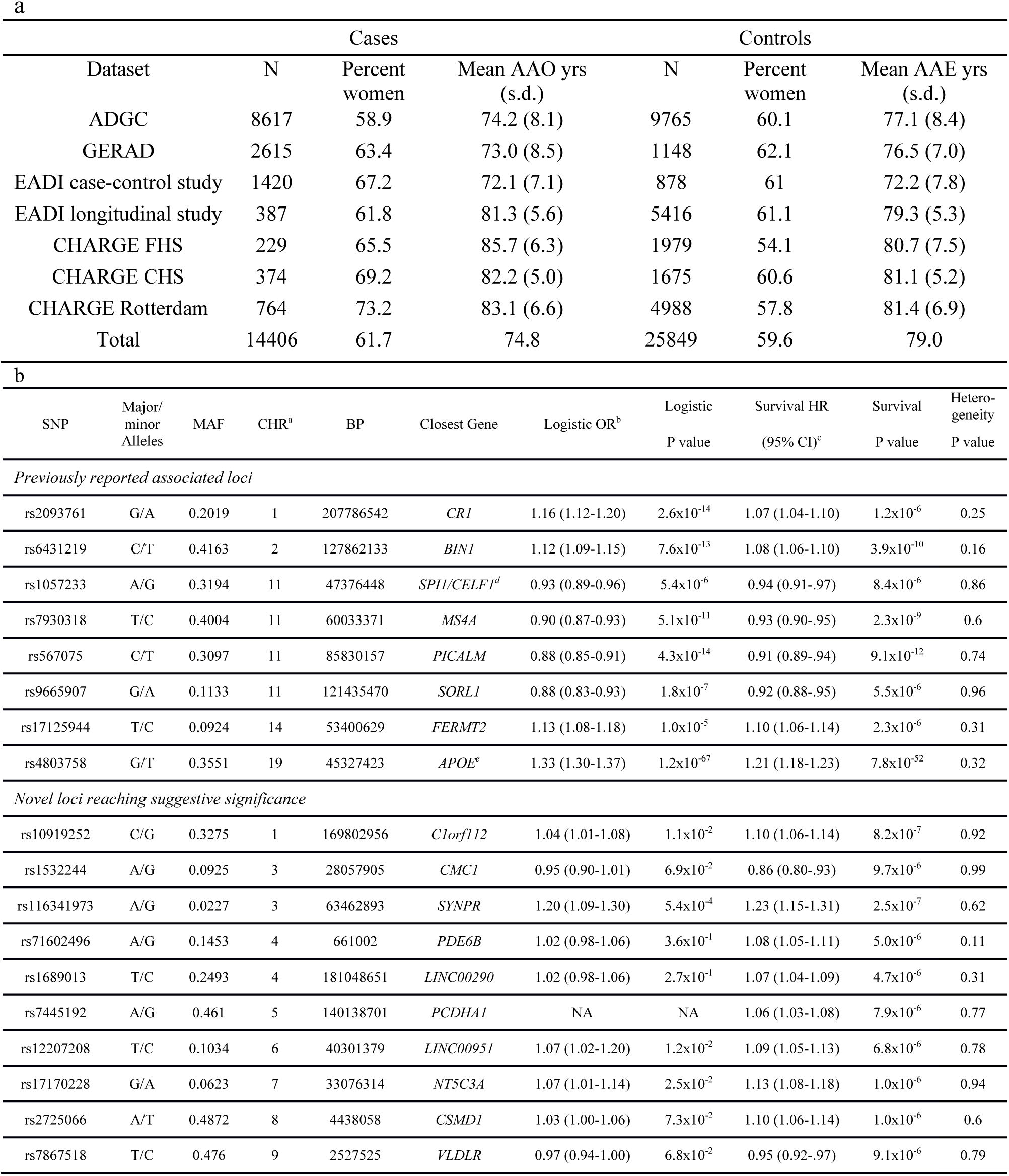
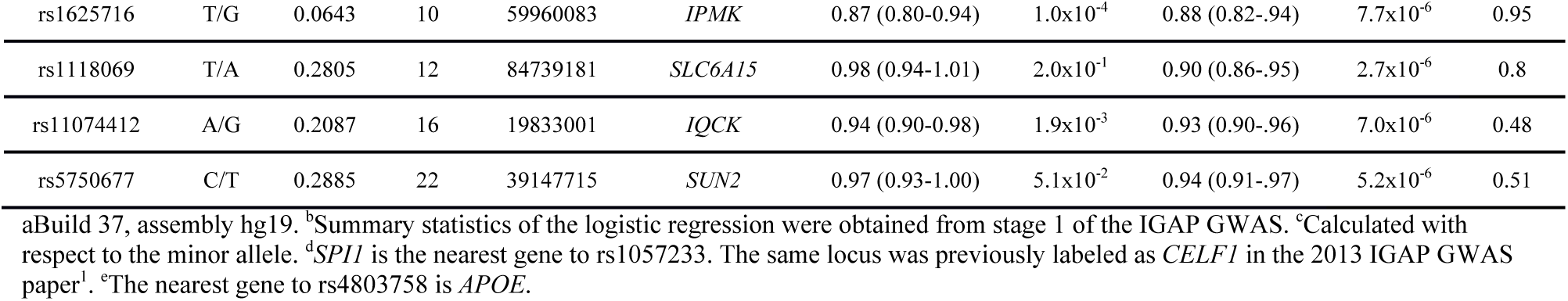
Genome-wide survival analysis of Alzheimer’s Disease. (a) Description of Consortia samples with available phenotype and genotype data included in the genome-wide survival analysis. AAO: age at onset. AAE: age at last examination. (b) Summary of loci with significant (P <5×10^−8^) or suggestive (P <1×10^−5^) associations from the genome-wide survival analysis.

### Cerebrospinal fluid biomarkers associations

To further validate the 22 loci with at least suggestive associations to AAOS, we examined their associations with CSF biomarkers, including total tau, phosphorylated tau_181_, and Aβ_42_ in a dataset of 3,646 Caucasians extended from our previous report^12^ (Table 2). Two SNPs showed associations that reached the Bonferroni-corrected threshold (P <2.27×10^−3^). Rs4803758 near *APOE* showed the most significant associations with levels of CSF phosphorylated tau_181_ (P=3.75×10^−4^) and CSF Aβ_42_ (P=3.12×10^−5^), whereas rs1057233 in the *SPI1/CELF1* locus was significantly associated with CSF Aβ_42_ (P=8.24×10^−4^). Of note, a SNP adjacent to *VLDLR*, rs7867518, showed the most significant association with CSF total tau (P=3.02×10^−3^), but failed to pass the Bonferroni-corrected threshold. The protective and deleterious effects in the survival analysis of these three SNPs were concordant with directionalities of their CSF biomarker associations; for example, the protective rs1057233^G^ allele was associated with higher CSF Aβ_42_ levels and the risk rs1057233^A^ allele was associated with lower CSF Aβ_42_ levels.

**Table 2.** Summary of CSF biomarker-associations of suggestive and significant AAOS-associated SNPs. Associations reaching the significance threshold after Bonferroni correction for multiple testing (P <2.27×10^−3^) are bolded.

| SNP | CHR | Closest gene | Beta<sub>tau</sub> | P<sub>tau</sub> | Beta<sub>ptau</sub> | P<sub>ptau</sub> | Beta<sub>ab42</sub> | P<sub>ab42</sub> |
| --- | --- | --- | --- | --- | --- | --- | --- | --- |
| *Previously reported associated loci* |  |  |  |  |  |  |  |  |
| rs2093761 | 1 | *CR1* | - | >0.05 | 1.46x10<sup>-2</sup> | 2.87x10<sup>-2</sup> | - | >0.05 |
| rs6431219 | 2 | *BIN1* | - | >0.05 | - | >0.05 | - | >0.05 |
| rs1057233 | 11 | *CELF1* | -1.11x10<sup>-2</sup> | 6.55x10<sup>-2</sup> | -1.25x10<sup>-2</sup> | 2.76x10<sup>-2</sup> | **1.45x10<sup>-2</sup>** | **8.24x10<sup>-4</sup>** |
| rs7930318 | 11 | *MS4A* | -1.24x10<sup>-2</sup> | 3.27x10<sup>-2</sup> | - | >0.05 | - | >0.05 |
| rs567075 | 11 | *PICALM* | -1.32x10<sup>-2</sup> | 3.22x10<sup>-2</sup> | -1.24x10<sup>-2</sup> | 3.13x10<sup>-2</sup> | 9.10x10<sup>-3</sup> | 3.88x10<sup>-2</sup> |
| rs9665907 | 11 | *SORL1* | -1.74x10<sup>-2</sup> | 4.28x10<sup>-2</sup> | -1.94x10<sup>-2</sup> | 1.57x10<sup>-2</sup> | - | >0.05 |
| rs17125944 | 14 | *FERMT2* | 2.50x10<sup>-2</sup> | 8.71x10<sup>-3</sup> | 2.09x10<sup>-2</sup> | 2.09x10<sup>-2</sup> | -1.79x10<sup>-2</sup> | 8.90x10<sup>-3</sup> |
| rs4803758 | 19 | *APOE* | 1.61x10<sup>-2</sup> | 7.42x10<sup>-3</sup> | **2.01x10<sup>-2</sup>** | **3.75x10<sup>-4</sup>** | **-1.79x10<sup>-2</sup>** | **3.12x10<sup>-5</sup>** |
| *Novel candidate loci* |  |  |  |  |  |  |  |  |
| rs10919252 | 1 | *C1orf112* | - | >0.05 | - | >0.05 | - | >0.05 |
| rs1532244 | 3 | *CMC1* | - | >0.05 | 2.41x10<sup>-2</sup> | 1.23x10<sup>-2</sup> | - | >0.05 |
| rs116341973 | 3 | *SYNPR* | - | >0.05 | - | >0.05 | - | >0.05 |
| rs71602496 | 4 | *PDE6B* | - | >0.05 | - | >0.05 | - | >0.05 |
| rs1689013 | 4 | *LINC00290* | - | >0.05 | - | >0.05 | - | >0.05 |
| rs7445192 | 5 | *PCDHA1* | - | >0.05 | 1.38x10<sup>-2</sup> | 9.98x10<sup>-3</sup> | - | >0.05 |
| rs12207208 | 6 | *LINC00951* | - | >0.05 | - | >0.05 | - | >0.05 |
| rs17170228 | 7 | *NT5C3A* | - | >0.05 | - | >0.05 | - | >0.05 |
| rs2725066 | 8 | *CSMD1* | 1.20x10<sup>-2</sup> | 4.53x10<sup>-2</sup> | - | >0.05 | - | >0.05 |
| rs7867518 | 9 | *VLDLR* | -1.58x10<sup>-2</sup> | 5.83x10<sup>-3</sup> | - | >0.05 | - | >0.05 |
| rs1625716 | 10 | *IPMK* | - | >0.05 | - | >0.05 | - | >0.05 |
| rs1118069 | 12 | *SLC6A15* | - | >0.05 | - | >0.05 | -1.07x10<sup>-2</sup> | 1.56x10<sup>-2</sup> |
| rs11074412 | 16 | *IQCK* | - | >0.05 | - | >0.05 | - | >0.05 |
| rs5750677 | 22 | *SUN2* | 1.30x10<sup>-2</sup> | 3.55x10<sup>-2</sup> | - | >0.05 | - | >0.05 |

### *Cis*-eQTL associations and colocalization analysis

Multiple disease-associated GWAS SNPs have been identified as *cis*-eQTLs of disease genes in disease-relevant tissues/cell types^23^. We investigated *cis*-eQTL effects of the 22 AAOS-associated SNPs and their tagging SNPs (R^2^ ≥ 0.8, listed in **Supplementary Table 3**) in the BRAINEAC dataset. We identified 4 significant associations (Bonferroni-corrected threshold: P=0.05/292,000 probes=1.7×10^−7^): rs1057233 was associated with *MTCH2* expression in the cerebellum (P=1.20×10^−9^); rs7445192 was associated with *SRA1* expression averaged across brain regions (P=7.0×10^−9^, 1.6×10^−7^ for two probes respectively), and rs2093761 was associated with *CR1/CR1L* expression in white matter (P=1.30×10^−7^, **Supplementary Table 4**). Further analysis using the GTEx dataset^24^ identified 50 unique, associated snp-gene pairs across 44 tissues, including 11 snp-gene pairs in various brain regions (**Supplementary Table 5**).

Recently, genetic and molecular evidence has implicated myeloid cells in the etiology of AD, including our finding that AD risk alleles are enriched for *cis*-eQTL effects in monocytes but not CD4+ T-lymphocytes^21^. To extend this finding and identify relevant cell types in AD, we used stratified LD score regression to estimate enrichment of AD heritability (measured by summary statistics from IGAP GWAS^1^) partitioned by 220 cell type–specific functional annotations as described by Finucane et al.^25^. We found a significant enrichment of AD heritability in hematopoietic cells of the myeloid and B-lymphoid lineage (e.g., 14.49 fold enrichment, P=3.49×10^−5^ in monocytes/CD14 enhancers/H3K4me1 and 12.33 fold enrichment, P=1.41×10^−6^ in B-cells/CD19 enhancers/H3K4me1). In contrast schizophrenia (SCZ) heritability was not enriched in hematopoietic cells (1.24 fold enrichment, P=0.53, as measured by summary statistics from the Psychiatric Genomics Consortium [PGC] GWAS^26^) but was significantly enriched in brain (18.61 fold enrichment, P=1.38×10^−4^ in fetal brain promoters/H3K4me3, **Supplementary Table 6**). These results suggest that myeloid cells specifically modulate AD susceptibility.

Based on these observations, we hypothesized that *cis*-eQTL effects of some AD-associated alleles may be specific to myeloid cells and thus not easily detectable in *cis*-eQTL datasets obtained from brain homogenates where myeloid cells (microglia and other brain-resident macrophages) represent a minor fraction of the tissue. Therefore, we analyzed *cis*-eQTL effects of the AAOS-associated SNPs and their tagging SNPs in human *cis*-eQTL datasets composed of 738 monocyte and 593 macrophage samples from the Cardiogenics consortium^27^. We identified 14 genes with *cis*-eQTLs significantly associated with these SNPs (Table 3). Notably, the protective rs1057233^G^ allele, located within the 3’ UTR of *SPI1*, was strongly associated with lower expression of *SPI1* in both monocytes (P=1.50×10^−105^) and macrophages (P=6.41×10^−87^) (Fig. 1a, 1b, 2a). This allele was also associated with lower expression of *MYBPC3* (monocytes: P=5.58×10^−23^; macrophages: P=4.99×10^−51^), higher expression of *CELF1* in monocytes (P=3.95×10^−8^) and lower *NUP160* expression in macrophages (P=5.35×10^−22^). Each of these genes lies within the *SPI1/CELF1* locus, suggesting complex regulation of gene expression in this region. Within the *MS4A* locus, which contains many gene family members, the minor allele (C) of rs7930318 was consistently associated with lower expression of *MS4A4A* in monocytes (P=8.20×10^−28^) and *MS4A6A* in monocytes (P=4.90×10^−23^) and macrophages (P=1.25×10^−9^, Fig. 1b). Among the novel AAOS-associated loci, rs5750677 was significantly associated with lower expression of *SUN2* in both monocytes (P=3.66×10^−58^) and macrophages (P=3.15×10^−36^), rs10919252 was associated with lower expression of *SELL* in monocytes (P=7.33×10^−35^), and rs1625716 was associated with lower expression of *CISD1* in macrophages (P=5.98×10^−23^, Table 3).

**Figure 1.**
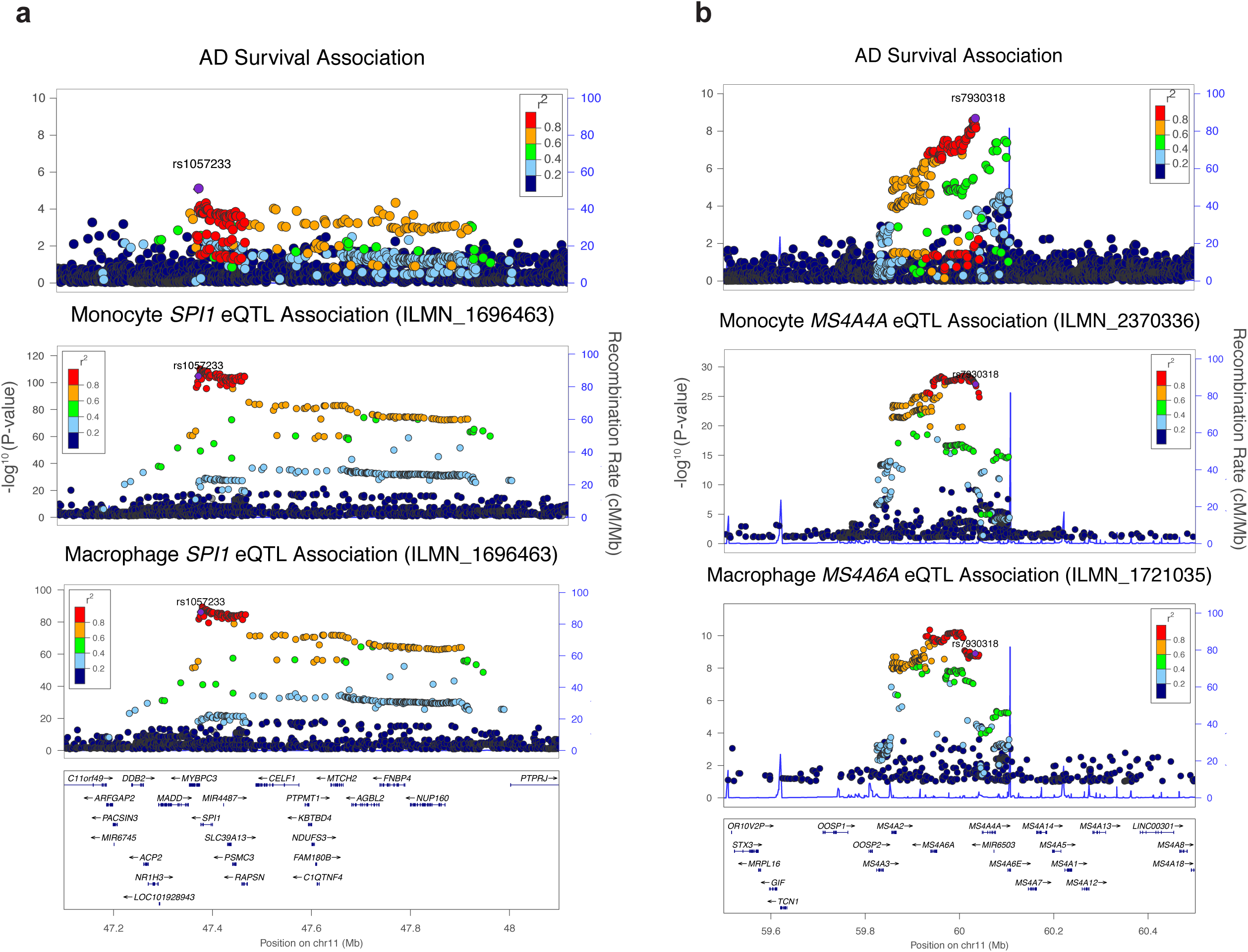
Genetic and eQTL fine-mapping of AD. (**a**) The AD-survival association landscape at the *CELF1/SPI1* locus resembles that of *SPI1* eQTL association in monocytes and macrophages. (**b**) The AD-survival association landscape resembles that of *MS4A4A/MS4A6A* eQTL association in monocytes and macrophages.

**Table 3.**
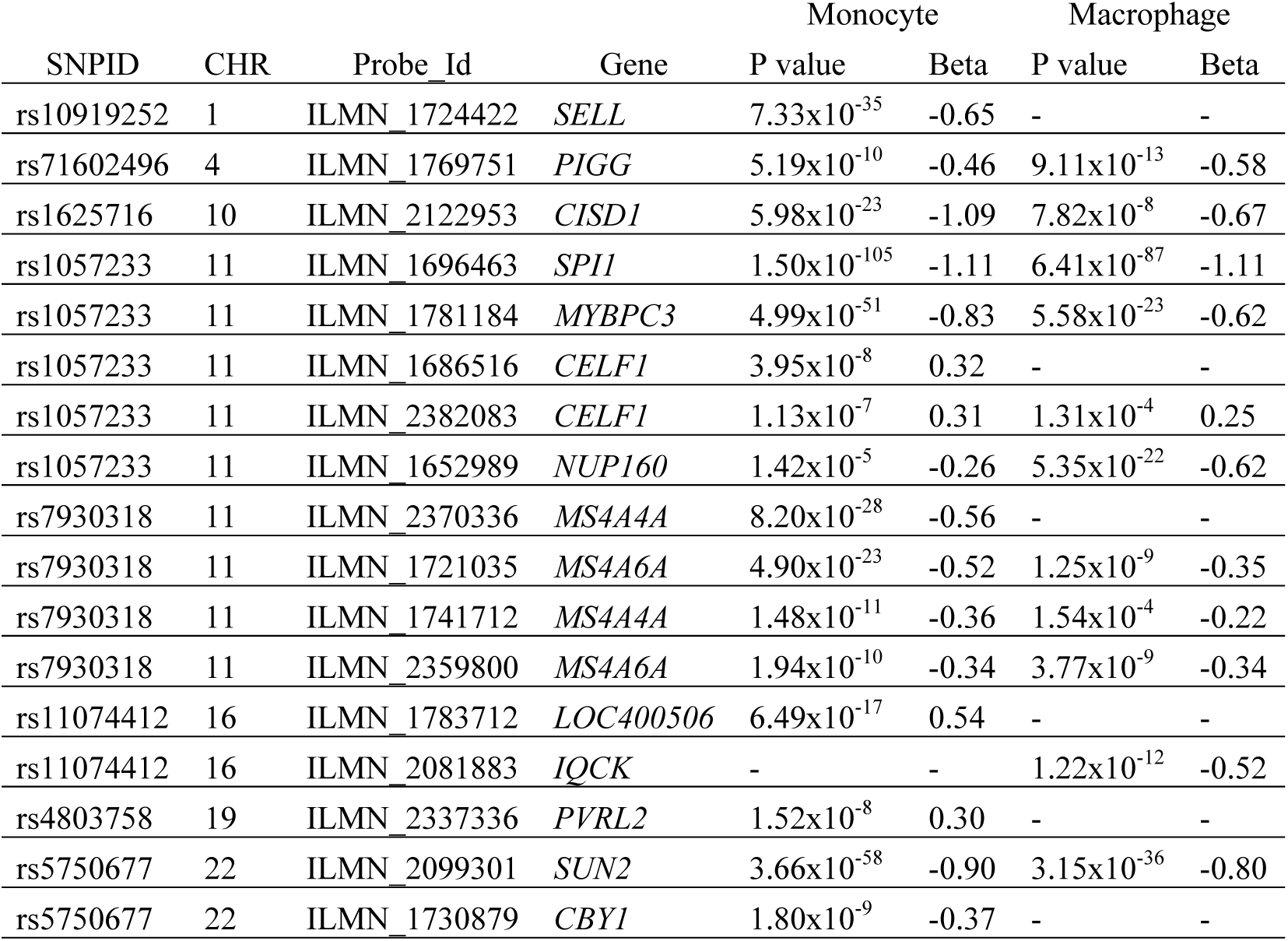
Significant *cis*-eQTL associations of the 22 suggestive and significant AAOS-associated SNPs. Significance threshold is determined to be 2.52×10^−6^ based on Bonferroni correction for multiple testing. The minor alleles are considered as the effective allele.

We then sought evidence of replication in an independent dataset of primary CD14+ human monocytes from 432 individuals^28^. We replicated *cis*-eQTL associations with expression of *SPI1*, *MYBPC3*, *MS4A4A*, *MS4A6A*, and *SELL* (Bonferroni-corrected threshold: P=0.05/15421 probes=3.24×10^−6^). We found strong evidence for the association between rs1057233 and *SPI1* expression (P=6.39×10^−102^) as well as *MYBPC3* expression (P=5.95×10^−33^, **Supplementary Table 7**). Rs1530914 and rs7929589, both in high LD with rs7930318 (R^2^=0.99 and 0.87, respectively), were associated with expression of *MS4A4A* and *MS4A6A* (P=3.60×10^−8^, 6.37×10^−15^), respectively. Finally, rs2272918, tagging rs10919252, was significantly associated with expression of *SELL* (P=8.43×10^−16^). Interestingly, the minor allele of all of these SNPs showed protective effects in both AD risk and survival analyses, as well as lower expression of the associated genes. Further, *SPI1*, *MS4A4A*, *MS4A6A*, and *SELL* are specifically expressed in microglia based on RNA-Seq data^29–31^ (Fig. 2b, Supplementary Fig. 4). However, *MYBPC3/*Mybpc3 (a myosin binding protein expressed at high levels in cardiac muscle cells) is either not expressed or expressed at low levels in human and mouse microglia, respectively. Amongst all genes probed, *MYBPC3* (ILMN_1781184) expression is the most highly and significantly correlated with *SPI1* (ILMN_1696463) expression in both Cardiogenics datasets (Spearman’s rho=0.54, qval=0.00 in monocytes and Spearman’s rho=0.42, qval=0.00 in macrophages) suggesting that low levels of *MYBPC3* expression in human myeloid cells are possibly due to leaky transcription driven by the adjacent highly expressed *SPI1* gene.

**Figure 2.**
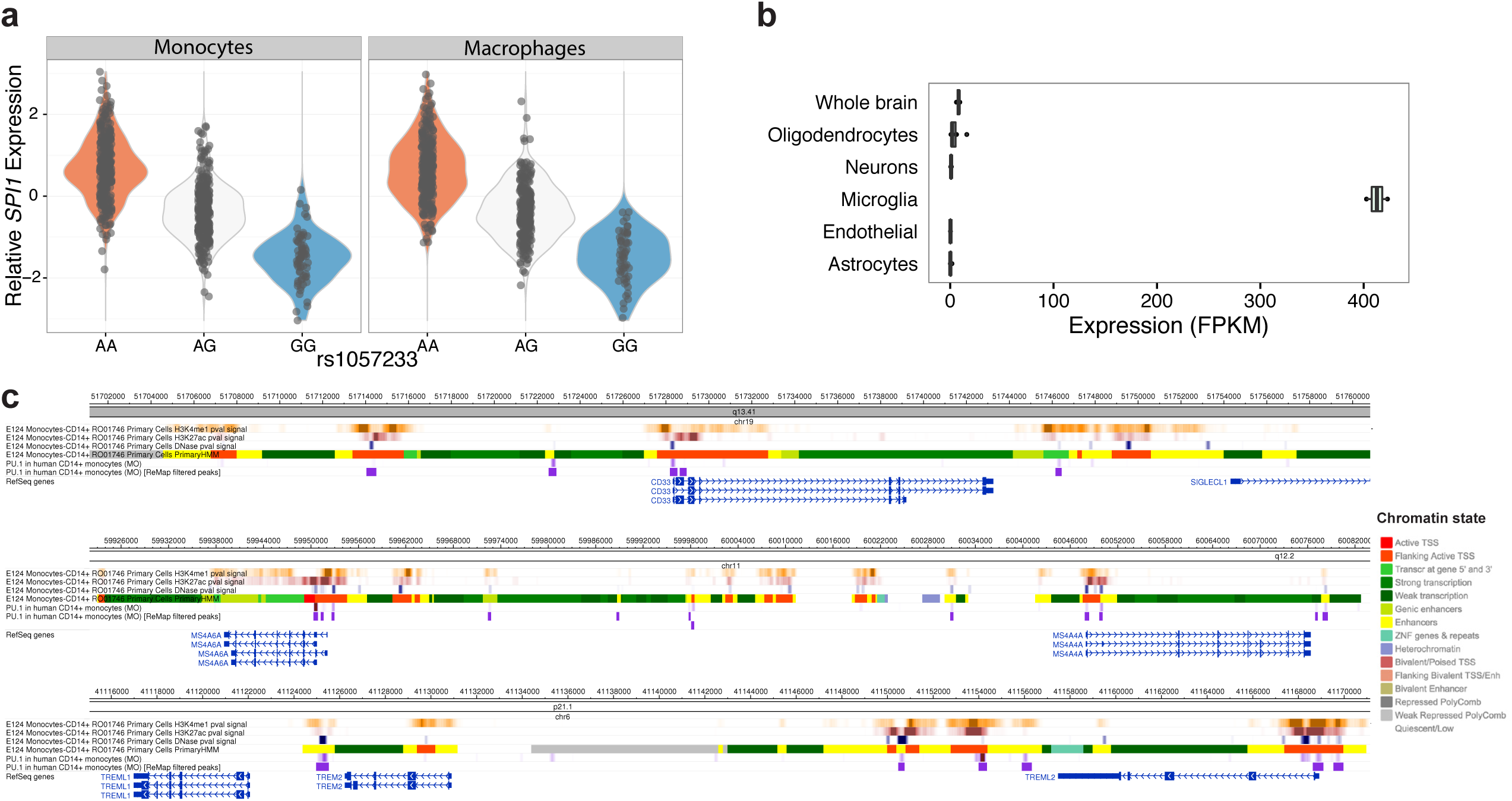
*SPI1* (PU.1) expression and ChIP-Seq analysis. (**a**) Rs1057233^G^ is associated with reduced *SPI1* expression in a dosage-dependent manner. (**b**) The mouse homolog of *SPI1*, *Sfpi1* or *Spi1*, is selectively expressed in microglia and macrophages in mouse brains based on the brain RNA-Seq database^29–31^. OPCs contain 5% microglial contamination. (**c**) *SPI1* (PU.1) binds to the promoter and regulatory regions of *CD33*, *MS4A4A*, *MS4A6A*, *TREM2*, and *TREML2* in human CD14+ monocytes based on ChIP-Seq data^35^.

We performed the coloc test^32^ to determine whether AAOS-associated SNPs co-localize with myeloid *cis*-eQTLs at the *SPI1/CELF1, MS4A* and *SELL* loci. These analyses (**Supplementary Table 8**) highlighted *SPI1* at the *SPI1/CELF1* locus as the strongest and most consistent colocalization target, and the only gene where the AD survival and gene expression association signals are likely (posterior probability ≥ 0.8) driven by the same causal genetic variant, in both monocytes and macrophages (PP.H4.abf of 0.85 and 0.83, respectively). *MYBPC3* in the *SPI1/CELF1* locus and *MS4A6A* in the *MS4A* locus also showed evidence of colocalization in both myeloid cell types albeit not surviving posterior probability cutoff in one of them. *MS4A4A* and *MS4A6E* in the *MS4A* locus showed evidence of co-localization only in monocytes, while *SELL* did not show evidence of colocalization.

In light of the strong *cis*-eQTL effects and colocalization results described above, we decided to focus subsequent analyses on *SPI1* as the strongest candidate gene underlying the disease association in myeloid cells.

### Conditional and SMR analysis of the *SPI1/CELF1* locus

The AAOS-association landscape shows that highly associated SNPs at the *SPI1/CELF1* locus span multiple genes (Fig. 1a). In the previous IGAP GWAS^1^, rs10838725 showed the strongest association at this locus (P=6.7×10^−6^, 1.1×10^−8^ vs. rs1057233: P=5.4×10^−6^, 5.9×10^−7^ in IGAP stage I, stage I and II combined, respectively). Rs10838725 is located in the intron of *CELF1*, which was assigned as the putative causal gene at this locus^1^ based on proximity to the index SNP, a criterion that has often proven to be erroneous^14^. In our survival analysis, however, rs10838725 showed weak association (P=0.12, HR=1.02, 95% CI=0.99-1.05) whereas rs1057233, located in the 3’UTR of a neighboring gene, *SPI1*, showed the strongest association (Table 1, P=5.4×10^−6^). The two SNPs exhibit only moderate linkage disequilibrium in the ADGC subset of the IGAP GWAS (R^2^=0.21, D’=0.96). Applying conditional logistic regression analysis of AD risk in the ADGC dataset, we found that rs1057233 remained significantly associated with AD after adjusting for rs10838725 (P=3.2×10^−4^), whereas rs10838725 showed no evidence of association after adjusting for rs1057233 (P=0.66). This suggests that rs1057233 is in stronger LD with the AD risk causal variant.

The association landscape in the AD survival analysis highly resembles that of *SPI1 cis*-eQTL analysis in myeloid cells (Fig. 1a). We reasoned that the associations of rs1057233 with AD-related phenotypes may be explained by the regulation of *SPI1* expression in myeloid cells, and that conditional analysis of the *cis*-eQTL signal could help us further dissect this complex locus. Therefore, we conducted conditional *cis*-eQTL analyses in both Cardiogenics datasets as we did above using rs1057233 (the top SNP for AD survival) and rs10838725 (the top SNP for AD risk). In addition, we also examined rs10838698 (a SNP in high LD with rs1057233 that was directly genotyped in the Cardiogenics dataset) and rs1377416, a SNP in high LD with rs10838725 proposed to be a functional variant in an enhancer near *SPI1* that is active in human myeloid cells and in the brain of a mouse model of AD^19^. It should be noted that rs1057233 is a functional variant that has been shown to directly affect *SPI1* expression by changing the target sequence and binding of miR-569^33^. Rs1057233 and rs10838698 remained significantly associated with *SPI1* expression when adjusting for either of the other two SNPs in both monocytes and macrophages (P <8.33×10^−3^). However, conditioning for either of these two SNPs abolished the associations of rs1377416 and rs10838725 with *SPI1* expression (**Supplementary Table 9**). Thus, the functional variant(s) mediating the effect on *SPI1* expression and AD risk likely reside(s) in the LD block that includes rs1057233 and rs10838698 but not rs10838725 and rs1377416 (Supplementary Fig. 5).

Using HaploReg^34^ to annotate the top AAOS-associated SNP (rs1057233) and its tagging SNPs (R^2^ >= 0.8, **Supplementary Table 3**), we identified multiple SNPs (e.g, rs10838699 and rs7928163) in tight LD with rs1057233 that changed the predicted DNA binding motif of SPI1 (PU.1), raising the possibility of altered self-regulation associated with the minor allele. Based on these results, one or more of these or other SNPs in very high LD with rs1057233, could explain the observed associations with *SPI1* expression and AD-related phenotypes.

We also conducted Summary-data-based Mendelian Randomization (SMR) and Heterogeneity In Dependent Instruments (HEIDI) tests^23^ to prioritize likely causal genes and variants by integrating summary statistics from our AAOS GWAS and the Cardiogenics study (**Supplementary Table 10**). SMR/HEIDI analysis was performed for the *SPI1/CELF1* locus using rs1057233, rs10838698, rs10838699, rs7928163, rs10838725 and rs1377416 as candidate causal variants. In both monocytes and macrophages, *SPI1* was consistently identified as the most likely gene whose expression levels are associated with AD survival because of causality/pleiotropy at the same underlying causal variant (rs1057233, rs10838698, rs10838699, or rs7928163 in the same LD block) (SMR P <4.90E-04, Bonferroni-corrected threshold for 6 SNPs tested against 17 probes and HEIDI P ≥ 0.05, Supplementary Fig. 6). Neither conditional analysis nor SMR/HEIDI analysis could definitively identify a single functional variant among this set of 4 SNPs in high LD. Functional analyses will be necessary to determine which SNPs in this LD block directly affects *SPI1* expression. Overall, rs1057233 and tagging SNPs are associated with AD risk and survival, and CSF Aβ_42_. The strong *cis*-eQTL effects and colocalization results point to *SPI1* as the most likely candidate gene underlying the disease association at the *SPI1*/*CELF1* locus.

### SPI1 (PU.1) cistrome and functional analysis in myeloid cells

*SPI1* encodes PU.1, a transcription factor essential for the development and function of myeloid cells. We hypothesize that it may modulate AD risk by regulating the transcription of AD-associated genes expressed in microglia and/or other myeloid cell types. First, we tested AD-associated genes for evidence of expression in human microglia^29^ as well as presence of PU.1 binding peaks in *cis*-regulatory elements of these genes using ChIP-Seq datasets obtained from human monocytes and macrophages^35^. We specifically investigated 112 AD-associated genes, including the 104 genes located within IGAP GWAS loci^36^ and *APOE, APP, TREM2* and *TREML2*, *TYROBP, TRIP4, CD33*, and *PLD3*. Among these genes, 75 had evidence of gene expression in human brain microglial cells, 60 of which also had evidence of association with PU.1 binding sites in human blood myeloid cells^35^ (**Supplementary Table 11**). Further examination of PU.1 binding peaks and chromatin marks/states in human monocytes confirmed that PU.1 is bound to *cis*-regulatory elements of many AD-associated genes, including *ABCA7*, *CD33*, *MS4A4A*, *MS4A6A*, *PILRA*, *PILRB, TREM2, TREML2*, and *TYROBP* (as well as *SPI1* itself, but notably not *APOE*) (Fig. 2c, Supplementary Fig. 7). Together, these results suggest that PU.1 may regulate the expression of multiple AD-associated genes in myeloid cells.

To further support that PU.1 target genes expressed in myeloid cells may be associated with AD risk, we used stratified LD score regression^25^ to estimate enrichment of AD heritability (as measured by summary statistics from the IGAP GWAS^1^) partitioned on the PU.1 cistrome, as profiled by ChIP-Seq in human monocytes and macrophages^35^. We found a significant enrichment of AD heritability in both monocytes (47.58 fold enrichment, P=6.94×10^−3^) and macrophages (53.88 fold enrichment, P=1.65×10^−3^), but not SCZ heritability [as measured by summary statistics from the PGC GWAS^26^] (**Supplementary Table 12**). Thus, the contribution of the myeloid PU.1 target gene network to disease susceptibility is specific to AD. However, since PU.1 is a key myeloid transcription factor that regulates the expression of a large number of genes in myeloid cells, the enrichment of AD risk alleles in PU.1 binding sites could simply reflect an enrichment of AD GWAS associations for genes that are expressed in myeloid cells rather than specifically among PU.1 target genes. To attempt to address this issue, we performed stratified LD score regression of AD heritability partitioned by functional annotations obtained from *SPI1* (marking the PU.1 cistrome) and POLR2AphosphoS5 (marking actively transcribed genes) ChIP-Seq experiments, performed in duplicate, using a human myeloid cell line (HL60) by the ENCODE Consortium^37^. We observed a significant enrichment for *SPI1* (PU.1) (34.58 fold enrichment, P=1.31×10^−3^ in first replicate; 58.12 fold enrichment, P=4.95×10^−3^ in second replicate) much stronger than that for POLR2AphosphoS5 (15.78 fold enrichment, P=1.71×10^−2^ in first replicate; 16.34 fold enrichment, P=1.25×10^−1^ in second replicate), consistent with our hypothesis (**Supplementary Table 12**).

PU.1 target genes are implicated in various biological processes of myeloid cells that may modulate AD risk. For example, a microglial gene network for pathogen phagocytosis has been previously implicated in the etiology of AD^18^. We modulated levels of PU.1 by *Spi1* cDNA overexpression or shRNA knock-down in BV2 mouse microglial cells, and used zymosan bioparticles labeled with pHrodo (a pH-sensitive dye that emits a fluorescent signal when internalized in acidic vesicles during phagocytosis) to measure pathogen engulfment. Analysis of zymosan uptake by flow cytometry revealed that phagocytic activity is augmented in BV2 cells overexpressing PU.1 (Fig. 3a), while knock-down of PU.1 resulted in decreased phagocytic activity (Fig. 3a). We confirmed overexpression and knock-down of PU.1 expression levels by western blotting and qPCR (Fig. 3). Phagocytic activity was not changed in untransfected cells when analyzed by flow cytometry (Supplementary Fig. 8d, 8e, 8f, 8g). These data suggest that modulation of PU.1 expression levels significant changes microglial phagocytic activity in response to fungal targets (mimicked by zymosan).

**Figure 3.**
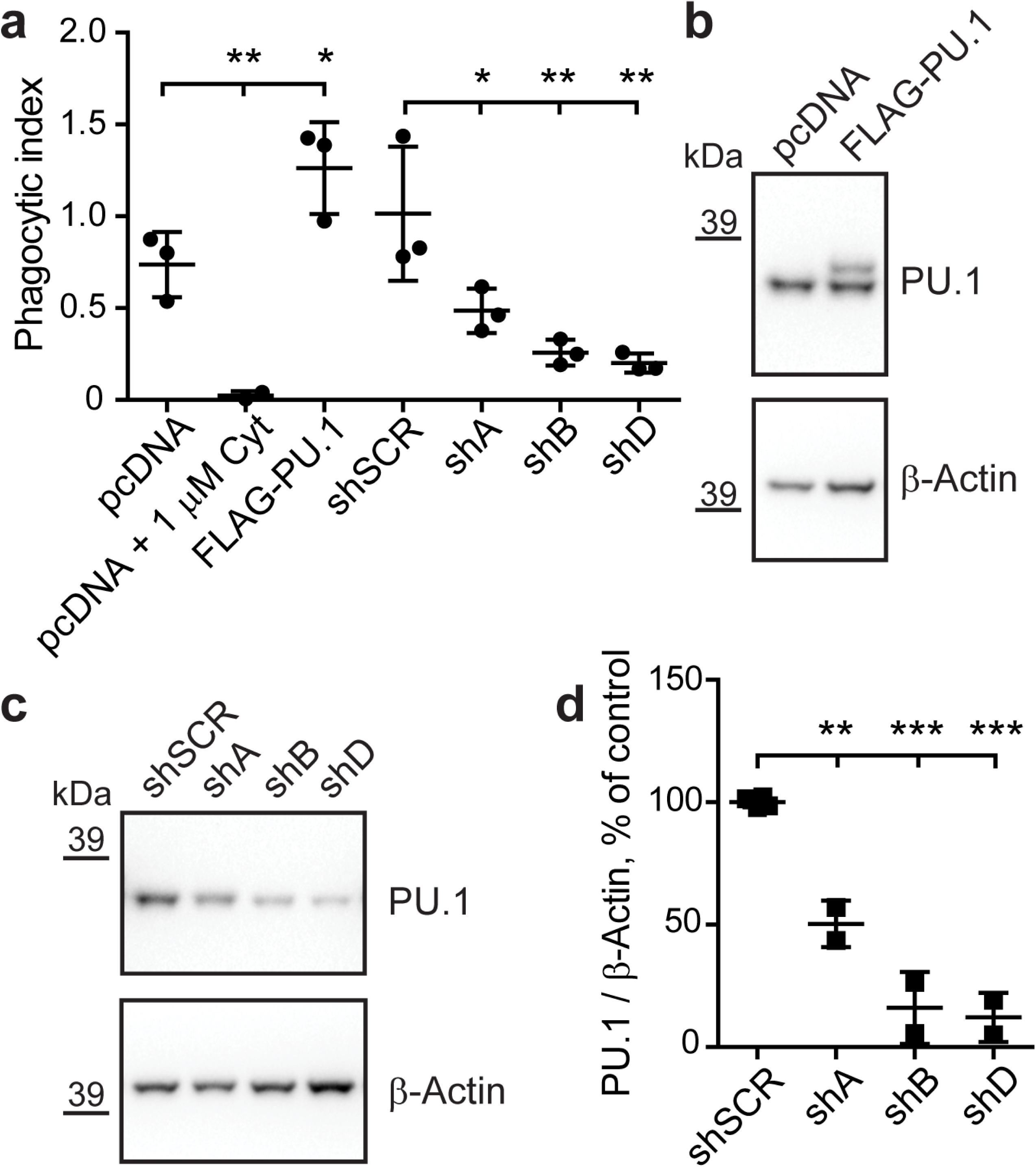
PU.1 modulates the phagocytic activity of BV2 microglial cells. (**a**) Phagocytosis of zymosan labeled with red pHrodo fluorescent dye in BV2 cells with transient overexpression and knock-down of PU.1 was measured by flow cytometry. Cytochalasin D treatment was used as a negative control. Mean phagocytic index ± SD is shown: pcDNA 0.7373 ± 0.1772, pcDNA + 1 μM Cyt 0.0236 ± 0.0242, FLAG-PU.1 1.2630 ± 0.2503, shSCR 1.014 ± 0.3656, shA 0.4854 ± 0.1209, shB 0.2579 ± 0.06967, shD 0.2002 ± 0.05168. F(6,13)=14.82, pcDNA vs pcDNA + 1 μM Cyt P=0.0078, pcDNA vs FLAG-PU.1 P=0.0295, shSCR vs shA P=0.0283, shSCR vs shB P=0.0020, shSCR vs shD P=0.0010, n=3. (**b**) BV2 cells were transiently transfected with pcDNA3 (pcDNA) or pcDNA3-FLAG-PU.1 (FLAG-PU.1) and pCMV-GFP as described for phagocytosis assay. Note a shift in mobility of the band for exogenous FLAG-PU.1 in overexpression condition compared to endogenous PU.1 in control. (**c**) BV2 cells were transiently transfected with shRNA targeting PU.1 (shA, shB and shD) or non-targeting control (shSCR) in pGFP-V-RS vector. GFP^+^ cells were sorted with flow cytometer and analyzed for levels of PU.1 in western blotting in two independent experiments (**b**, **c**). (**d**) Quantification of PU.1 levels in **c** normalized to β-Actin as a loading control. Values are presented as mean ± SD: shSCR 100 ± 2.10, shA 50.34 ± 9.52, shB 16.03 ± 14.72, shD 12.13 ± 10.03. F(3,6)=70.55, shSCR vs shA P=0.0014, shSCR vs shB P <0.0001, shSCR vs shD P <0.0001, n=2. * P <0.05, ** P <0.01, *** P <0.001, one-way ANOVA with Sidak’s post hoc multiple comparisons test between selected groups.

To further explore the functional impact of variation in *SPI1* expression, we performed qPCR to test whether differential *Spi1* expression in BV2 cells can modulate expression of genes thought to play important roles in AD pathogenesis and/or microglial cell function (Fig. 3, Supplementary Fig. 9, **Supplementary Table 13, 14**). We found that levels of some of these genes were affected in opposing directions by overexpression and knock-down of *Spi1* (Fig. 4a), while other genes were affected only by overexpression (Fig. 4b) or knock-down (Fig. 4c) or not affected at all (Supplementary Fig. 9). After knock-down of *Spi1* in BV2 cells, expression of *Cd33*, *Tyrobp*, *Ms4a4a* and *Ms4a6d* decreased and expression of *Apoe* and *Clu/ApoJ* increased (Fig. 4a, 4c). These data demonstrate that multiple microglial genes, some already implicated in AD, are selectively perturbed by altered expression of *Spi1*.

**Figure 4.**
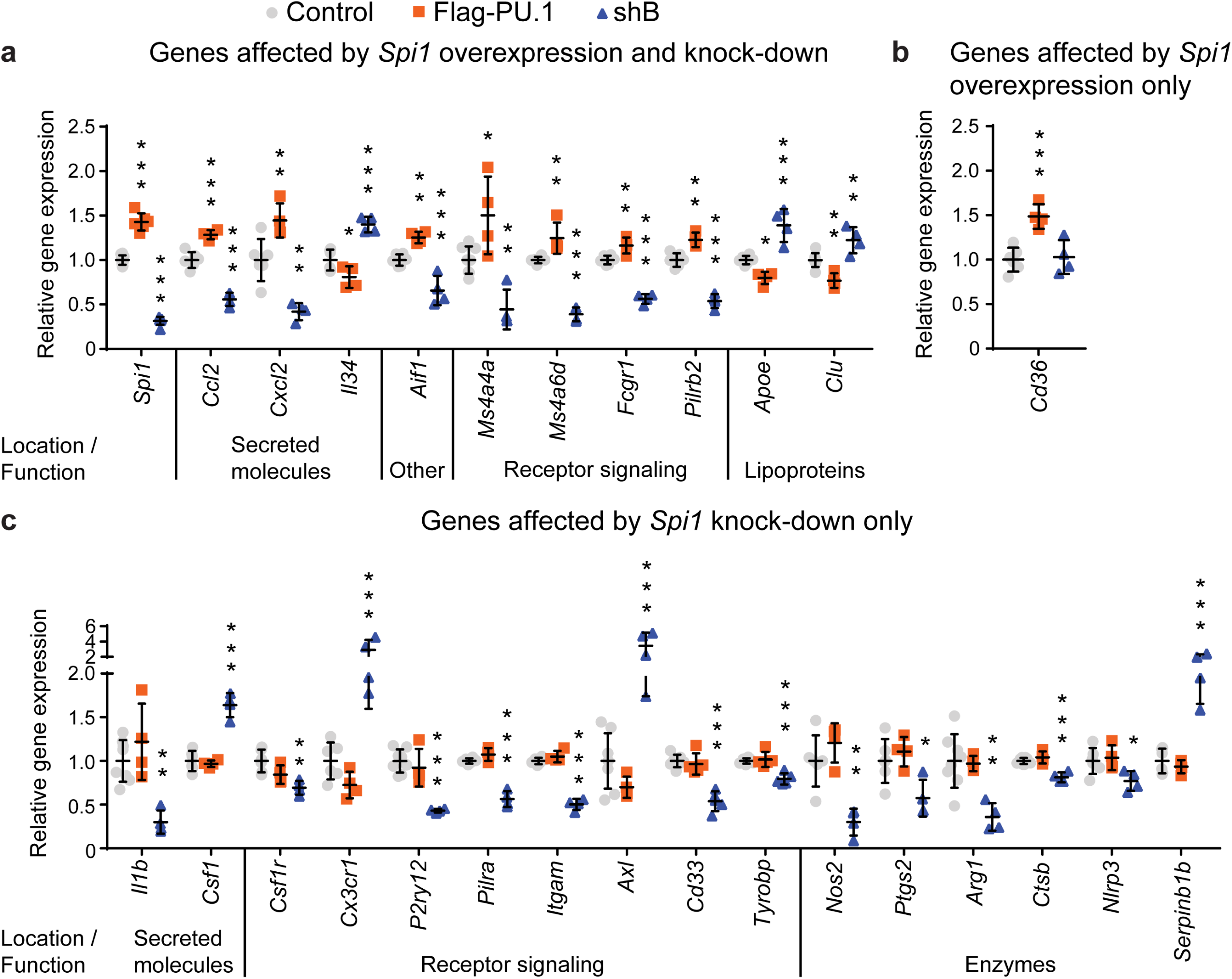
Genes regulated with differential expression of *Spi1* in BV2 microglial cells. qPCR analysis in transiently transfected and sorted GFP^+^ BV2 cells with overexpression (FLAG-PU.1) and knock-down (shB) of *Spi1*. Changes in expression levels are grouped for genes with altered levels after overexpression and knock-down of *Spi1* in (**a**) and genes with variable expression in BV2 cells either with overexpression (**b**) or knock-down (**c**) of *Spi1*. Values are presented as mean ± SD, n=4 samples collected independently. * P <0.05, ** P <0.01, *** P <0.001, one-way ANOVA with Dunnett’s post hoc multiple comparisons test between experimental and control groups, detailed statistical analysis is reported in **Supplementary Table 11**.

## Discussion

By performing a large-scale genome-wide survival analysis, we discovered multiple loci associated with AAOS (Table 1). The four genome-wide significantly associated loci, *BIN1* (P=7.6×10^−13^), *MS4A* (P=5.1×10^−11^), *PICALM* (P=4.3×10^−14^), and *APOE* (P=1.2×10^−67^), have been previously reported to be associated with AD risk^1^. Notably, this is the first study showing that the *MS4A* locus is associated with AAOS. The most significantly AAOS-associated SNP at this locus, rs7930318, shows a protective effect (HR=0.93, 95% CI=0.90−.95) in the survival analysis, consistent with the previous IGAP GWAS logistic regression analysis for AD risk (OR=0.90, 95% CI=0.87-.93).

By combining AAOS and CSF biomarker GWAS results, we provide evidence of AD association at additional loci (Table 2). In particular, rs7867518 at the *VLDLR* locus shows suggestive associations with both AAOS (P=9.1×10^−6^) and CSF tau (P=3.03×10^−3^). An adjacent SNP (rs2034764) in the neighboring gene, *KCNV2*, has been previously reported to have suggestive association with AAO^22^. VLDLR is a receptor for lipoproteins containing APOE^38^ and CLU/APOJ^39^, another AD risk gene. Additionally, the *VLDLR*-5-repeat allele was found to be associated with dementia^38^. This genetic and biochemical evidence suggests *VLDLR* may be linked to AD.

*Cis*-eQTL analyses of AAOS-associated SNPs revealed limited associations when using data from brain tissue homogenates, yet identified multiple candidate genes when using data from myeloid cells, the top candidate causal cell types for AD based on the stratified LD score regression analysis of AD heritability presented here. This calls attention to careful selection of relevant cell types in eQTL studies of disease associations. In particular, by conducting *cis*-eQTL analyses using monocyte and macrophage datasets, we discovered associations of AAOS-associated SNPs with the expression of *SELL*, *SPI1*, *MYBPC3*, *NUP160*, *MS4A4A*, *MS4A6A* and *SUN2* (Table 3). Furthermore, we replicated the *cis*-eQTL associations of rs1057233 with *SPI1, MYBPC3*, rs7930318 with *MS4A4A*, *MS4A6A* and rs2272918 with *SELL* in an independent monocyte dataset. We further showed that *SPI1* myeloid *cis*-eQTLs and AAOS-associated SNPs are not likely to be colocalized by chance and thus may be in the causal path to AD (Fig. 1). Notably, the minor allele of rs1057233 (G) is suggestively associated with lower AD risk (P=5.4×10^−6^, 5.9×10^−7^ in IGAP stage I, stage I and II combined, respectively)^1^, later AAO (P=8.4×10^−6^) and significantly associated with higher CSF Aβ_42_ (P=4.11×10^−4^), which likely reflects decreased Aβ aggregation and β-amyloid deposition in the brain. Furthermore, it is strongly associated with lower *SPI1* expression in human monocytes (P=1.50×10^−105^) and macrophages (P=6.41×10^−87^, Table 3).

Colocalization analyses using coloc^32^ and SMR/HEIDI^23^ support the hypothesis that the same causal SNP(s) influence *SPI1* expression and AD risk. However, neither conditional nor SMR/HEIDI analyses were able to pin-point an individual SNP; both approaches identified an LD block tagged by rs1057233, in which one or more SNPs may individually or in combination influence both *SPI1* expression and AD risk. rs1057233 changes the target sequence and binding of miR-569^33^, and its tagging SNPs alter binding motifs of transcription factors including PU.1 itself (**Supplementary Table 3** and Supplementary Fig. 7d). rs1377416, is located in a predicted enhancer in the vicinity of *SPI1* and altered enhancer activity when assayed *in vitro* using a reporter construct transfected in BV2 cells^19^. However, rs1057233 remained significantly associated with AD after conditioning for either rs1377416 (P=1.2×10^−3^) or the previously reported IGAP GWAS top SNP rs10838725 (P=3.2×10^−4^) in the ADGC dataset. Further, the *cis*-eQTL association between rs1057233 and *SPI1* expression remained significant after conditioning for either of these SNPs, whereas conditioning for rs1057233 abolished their *cis*-eQTL associations with *SPI1* (**Supplementary Table 9**). Thus, rs1057233 and its tagging SNPs likely represent the underlying disease locus and may modulate AD risk through variation in *SPI1* expression. Interestingly, rs1057233 was previously found to be associated with systemic lupus erythematosus^33^, body mass index^40^ and proinsulin levels^41^ and may contribute to the connection between AD, immune cell dysfunction, obesity and diabetes.

PU.1 binds to *cis*-regulatory elements of several AD-associated genes expressed in human myeloid cells, including *ABCA7*, *CD33, MS4A4A*, *MS4A6A*, *TREM2*, and *TYROBP* (Fig. 1e, Supplementary Fig. 7). Further, PU.1 binds to active enhancers of *Trem2* and *Tyrobp* in ChIP-Seq experiments using mouse BV2 cells^42^ or bone marrow-derived mouse macrophages^43^. PU.1 is required for the development and function of myeloid and B-lymphoid cells^44,45^. In particular, PU.1 expression is dynamically and tightly controlled during haematopoiesis to direct the specification of CD34+ hematopoietic stem and progenitor cells toward the myeloid and B-lymphoid lineage by progressively partitioning into CD14+ monocytes/macrophages, CD15+ neutrophils, and CD19+ B cells^46^, which are the cell types highlighted by our stratified LD score regression analysis. Given its selective expression in microglia in the brain (Fig. 2b), PU.1 may modify microglial cell function through transcriptional regulation of target genes that act as downstream modulators of AD susceptibility, as evidenced by the significant enrichment of AD heritability partitioned on the PU.1 cistrome in human myeloid cells (**Supplementary Table 12**).

In support of this hypothesis, we also demonstrate that changes in PU.1 expression levels alter phagocytic activity in BV2 mouse microglial cells (Fig. 3, Supplementary Fig. 8). Knock-down of PU.1 expression reduced engulfment of zymosan, whereas overexpression of PU.1 increased engulfment of zymosan, a Toll-like receptor 2 (TLR2) agonist that mimics fungal pathogens. This is in line with previous data showing decreased uptake of Aβ_42_ (also a TLR2 agonist) in primary microglial cells isolated from adult human brain tissue and transfected with siRNA targeting *SPI1*^47^. Interestingly, several AD-associated genes (e.g., *CD33*, *TYROBP*, *TREM2*, *TREML2*, *CR1*, *ABCA7*, *APOE*, *CLU/APOJ*) have been shown to be involved in the phagocytic clearance of pathogens or host-derived cellular material (e.g., β-amyloid, apoptotic cells, myelin debris, lipoproteins, etc.), suggesting a strong link between perturbation of microglial phagocytosis and AD pathogenesis. In addition to *Cd33, Tyrobp, Apoe* and *Clu/ApoJ*, several genes with roles in phagocytosis are dysregulated by altering *Spi1* expression, i.e. *Cd36*, *Fcgr1*, *P2ry12*, *Itgam*, *Cx3cr1*, *Axl*, *Ctsb* (Fig. 4a, 4b, 4c), suggesting a collective and coordinated effect of *Spi1* on the phagocytic activity of BV2 cells.

Our genetic analyses show that the protective allele at the *MS4A* locus is associated with lower expression of *MS4A4A* and *MS4A6A* in human myeloid cells, and the BV2 experiment demonstrated that lower expression of *Spi1* (which is protective in humans) led to lower expression of *Ms4a4a* and *Ms4a6d* (mouse ortholog of *MS4A6A*), which are also associated with reduced AD risk in humans. Transcriptomic and proteomic analyses of microglial cells suggested a microglial homeostatic signature that is perturbed during aging and under pathological conditions^48^. It will be valuable to test whether genetically altered *SPI1* levels prime microglia to exacerbate or alleviate transcriptional responses that occur during aging or disease development. Together with genetic variation in myeloid genes associated with AD as an amplifier, *SPI1* may be a master regulator capable of tipping the balance toward a neuroprotective or neurotoxic microglial phenotype.

PU.1 expression levels regulate multiple myeloid/microglial cell functions^47^, including proliferation, survival and differentiation, that could also modulate AD risk. Indeed, expression of *Il34* and *Csf1*, soluble factors that bind to *Csf1r* and required for microglial development and maintenance *in vivo*^49^, were elevated after knock-down of *Spi1*, while expression of *Csf1r* was reduced (Fig. 4a, 4c). Interestingly, inhibition of *Csf1r* in a 3xTg-AD mouse model led to a reduction in the number of microglia associated with β-amyloid plaques and improved cognition^50^. These findings suggest the importance of analyzing cell proliferation, survival, differentiation, and migration phenotypes with differential *Spi1* expression, because *Spi1* levels modulate expression of *Ccl2* and *Cxcl2* (Fig. 4a), which are MCP1 and MIP2α proteins that help recruitcirculating monocytes and neutrophils to the brain to promote neuroinflammation. In addition, knocking down *Spi1* reduced expression of a microgliosis marker *Aif1* (Iba1) along with *Il1b*, *Nos2*, *Ptgs2*, *Arg1* and *Nlrp3* (Fig. 4a, 4c), suggesting that decreased *Spi1* expression may blunt the pro-inflammatory response of microglial cells to improve disease outcomes.

Interestingly, expression of *Cx3cr1* and *Axl* were elevated upon knock-down of *Spi1* (Fig. 4c), raising the possibility that beneficial effects of changes in *Spi1* expression are exerted through modulation of synaptic or neuronal clearance. Further experimental investigation of these phenotypes may shed light on the mechanisms of *SPI1* modulation of AD risk. Of note, overexpression and knock-down of *Spi1* in BV2 cells produce different and often opposite changes in expression of the genes profiled here, possibly driving alternative phenotypes that may underlie detrimental and protective roles of PU.1.

In summary, by combining AD survival and endophenotype GWAS analyses, we replicated and discovered multiple genetic loci associated with AAOAAOS. Specifically, we nominate *SPI1* as the gene responsible for disease association at the previously reported *CELF1* locus. *SPI1* encodes PU.1, a transcription factor expressed in microglia and other myeloid cells that directly regulates other AD-associated genes expressed in these cell types. Our data suggest that lower *SPI1* expression reduces risk for AD, suggesting a novel therapeutic approach to the treatment of AD. Furthermore, we demonstrate that AAOS-associated SNPs within the MS4A gene cluster are associated with eQTLs in myeloid cells for both *MS4A4A* and *MS4A6A.* Specifically, the allele associated with reduced AD risk is associated with lower *MS4A4A* and *MS4A6A* expression. This is consistent with the observation that lowering *SPI1* expression, which is protective for AD risk, also lowers *MS4A4A* and *MS4A6A* expression. These results reinforce the emerging genetic and epigenetic association between AD and a network of microglial expressed genes^2,5,17–21^, highlighting the need to dissect their functional mechanisms.

## Acknowledgements

We would like to thank the patients, control subjects, and their family members for participating in or supporting the research projects included in this study. We thank Marc Diamond (UT Southwestern Medical Center) for the BV2 cell line and Flow Cytometry CORE at the Icahn School of Medicine at Mount Sinai Hospital.

## IGAP

GERAD was supported by the Wellcome Trust, the MRC, Alzheimer’s Research UK (ARUK) and the Welsh government. ADGC and CHARGE were supported by the US National Institutes of Health, National Institute on Aging (NIH-NIA), including grants U01 AG032984 and R01 AG033193. CHARGE was also supported by Erasmus Medical Center and Erasmus University.

## ADNI

Data collection and sharing for this project was funded by the Alzheimer’s Disease Neuroimaging Initiative (ADNI) (National Institutes of Health Grant U01 AG024904) and DOD ADNI (Department of Defense award number W81XWH-12-2-0012). ADNI is funded by the National Institute on Aging, the National Institute of Biomedical Imaging and Bioengineering, and through generous contributions from the following: AbbVie, Alzheimer’s Association; Alzheimer’s Drug Discovery Foundation; Araclon Biotech; BioClinica, Inc.; Biogen; Bristol-Myers Squibb Company; CereSpir, Inc.; Eisai Inc.; Elan Pharmaceuticals, Inc.; Eli Lilly and Company; EuroImmun; F. Hoffmann-La Roche Ltd and its affiliated company Genentech, Inc.; Fujirebio; GE Healthcare; IXICO Ltd.; Janssen Alzheimer Immunotherapy Research & Development, LLC.; Johnson & Johnson Pharmaceutical Research & Development LLC.; Lumosity; Lundbeck; Merck & Co., Inc.; Meso Scale Diagnostics, LLC.; NeuroRx Research; Neurotrack Technologies; Novartis Pharmaceuticals Corporation; Pfizer Inc.; Piramal Imaging; Servier; Takeda Pharmaceutical Company; and Transition Therapeutics. The Canadian Institutes of Health Research is providing funds to support ADNI clinical sites in Canada. Private sector contributions are facilitated by the Foundation for the National Institutes of Health (www.fnih.org). The grantee organization is the Northern California Institute for Research and Education, and the study is coordinated by the Alzheimer’s Disease Cooperative Study at the University of California, San Diego. ADNI data are disseminated by the Laboratory for Neuro Imaging at the University of Southern California

We thank the Cardiogenics (European Project reference LSHM-CT-2006-037593) project for providing summary statistics for the *cis*-eQTL-based analyses. We also thank the ENCODE Consortium and Richard Myers’ lab (HAIB) for providing ChIP-Seq datasets.

This work was supported by grants from the National Institutes of Health (U01 AG049508 (AMG), R01-AG044546 (CC), RF1AG053303 (CC) and R01-AG035083 (AMG), RF-AG054011 (AMG)), the JPB Foundation (AMG) and FBRI (AMG). The recruitment and clinical characterization of research participants at Washington University were supported by NIH P50 AG05681, P01 AG03991, and P01 AG026276. Kuan-lin Huang received fellowship funding in part from the Ministry of Education in Taiwan and the Lucille P. Markey Special Emphasis Pathway in Human Pathobiology. Ke Hao is partially supported by the National Natural Science Foundation of China (Grant No. 21477087, 91643201) and by the Ministry of Science and Technology of China (Grant No. 2016YFC0206507). This work was supported by access to equipment made possible by the Hope Center for Neurological Disorders and the Departments of Neurology and Psychiatry at Washington University School of Medicine.

## Author Contributions

A.M.G., E.M., and K.H. conceived and designed the experiments. K.H., S.C.J., O.H., A.D., M.K., J.C., J.C.L., V.C., C.B., B.G., Y.D., A.M., T.R., A.R., J.L.D., M.V.F, L.I., B.Z., I.B., C.C. and E.M. performed data analysis. A.A.P. performed phagocytosis assays, western blotting and qPCR validation. S.B., B.P.F., J.B., R.S., V.E.P., R.M., J.L.H., L.A.F., M.A.P., S.S., J.W., P.A., G.D.S., J.S.K.K., K.H., and C.C. provided and processed the data. A.M.G. supervised data analysis and functional experiments. K.H., A.A.P., E.M., and A.M.G. wrote and edited the manuscript. All authors read and approved the manuscript.

## Online Methods

### Genome-wide survival association study datasets

The final meta-analysis dataset consists of samples from the Alzheimer’s Disease Genetics Consortium (ADGC), Genetic and Environmental Risk in Alzheimer’s Disease (GERAD), European Alzheimer’s Disease Initiative (EADI), and Cohorts for Heart and Aging Research in Genomic Epidemiology (CHARGE). The study cohorts consist of case-control and longitudinal cohorts. For all studies, written informed consent was obtained from study participants or, for those with substantial cognitive impairment, from a caregiver, legal guardian, or other proxy, and the study protocols for all populations were reviewed and approved by the appropriate Institutional review boards. Details of ascertainment and diagnostic procedures for each dataset extend from details previously described^1–5^ and are documented below:

#### (1) Alzheimer’s Disease Genetics Consortium (ADGC)

The imputed ADGC sample that passed quality control procedures comprised of 8,617 AD cases and 9,765 control subjects from GWAS datasets assembled by the Alzheimer’s Disease Genetics Consortium (ADGC). Details of ascertainment and diagnostic procedures for each data set were as previously described^2^.

#### (2) Genetic and Environmental Risk in Alzheimer’s Disease (GERAD)

Data used in the preparation of this article were obtained from the Genetic and Environmental Risk for Alzheimer’s disease (GERAD) Consortium. The imputed GERAD sample comprised 3,177 AD cases and 7,277 controls with available age and gender data. A subset of this sample has been used in this study, comprising 2,615 cases and 1,148 elderly screened controls. Cases and elderly screened controls were recruited by the Medical Research Council (MRC) Genetic Resource for AD (Cardiff University; Institute of Psychiatry, London; Cambridge University; Trinity College Dublin), the Alzheimer’s Research UK (ARUK) Collaboration (University of Nottingham; University of Manchester; University of Southampton; University of Bristol; Queen’s University Belfast; the Oxford Project to Investigate Memory and Ageing (OPTIMA), Oxford University); Washington University, St Louis, United States; MRC PRION Unit, University College London; London and the South East Region AD project (LASER-AD), University College London; Competence Network of Dementia (CND) and Department of Psychiatry, University of Bonn, Germany; the National Institute of Mental Health (NIMH)AD Genetics Initiative. 6,129 population controls were drawn from large existing cohorts with available GWAS data, including the 1958 British Birth Cohort (1958BC) (http://www.b58cgene.sgul.ac.uk), the KORA F4 Study and the Heinz Nixdorf Recall Study. All AD cases met criteria for either probable (NINCDS-ADRDA, DSM-IV) or definite (CERAD) AD. All elderly controls were screened for dementia using the MMSE or ADAS-cog, were determined to be free from dementia at neuropathological examination or had a Braak score of 2.5 or lower. Genotypes from all cases and 4,617 controls were previously included in the AD GWAS by Harold and colleagues (2009). Genotypes for the remaining 2,660 population controls were obtained from WTCCC2. Imputation of the dataset was performed using IMPUTE2 and the 1000 genomes (http://www.1000genomes.org/) Dec2010 reference panel (NCBI build 37.1). The imputed data was then analysed using logistic regression including covariates for country of origin, gender, age and 3 principal components obtained with EIGENSTRAT software based on individual genotypes for the GERAD study participants.

GERAD Supplementary Acknowledgements: This study incorporated imputed summary results from the GERAD1 genome-wide association study. GERAD Acknowledgements: Cardiff University was supported by the Wellcome Trust, Medical Research Council (MRC), Alzheimer’s Research UK (ARUK) and the Welsh Assembly Government. Cambridge University and Kings College London acknowledge support from the MRC. ARUK supported sample collections at the South West Dementia Bank and the Universities of Nottingham, Manchester and Belfast. The Belfast group acknowledges support from the Alzheimer’s Society, Ulster Garden Villages, N.Ireland R&D Office and the Royal College of Physicians/Dunhill Medical Trust. The MRC and Mercer’s Institute for Research on Ageing supported the Trinity College group. The South West Dementia Brain Bank acknowledges support from Bristol Research into Alzheimer’s and Care of the Elderly. The Charles Wolfson Charitable Trust supported the OPTIMA group. Washington University was funded by NIH grants, Barnes Jewish Foundation and the Charles and Joanne Knight Alzheimer’s Research Initiative. Patient recruitment for the MRC Prion Unit/UCL Department of Neurodegenerative Disease collection was supported by the UCLH/UCL Biomedical Centre and NIHR Queen Square Dementia Biomedical Research Unit. LASER-AD was funded by Lundbeck SA. The Bonn group was supported by the German Federal Ministry of Education and Research (BMBF), Competence Network Dementia and Competence Network Degenerative Dementia, and by the Alfried Krupp von Bohlen und Halbach-Stiftung. The GERAD Consortium also used samples ascertained by the NIMH AD Genetics Initiative.

The KORA F4 studies were financed by Helmholtz Zentrum München; German Research Center for Environmental Health; BMBF; German National Genome Research Network and the Munich Center of Health Sciences. The Heinz Nixdorf Recall cohort was funded by the Heinz Nixdorf Foundation (Dr. jur. G.Schmidt, Chairman) and BMBF. Coriell Cell Repositories is supported by NINDS and the Intramural Research Program of the National Institute on Aging. We acknowledge use of genotype data from the 1958 Birth Cohort collection, funded by the MRC and the Wellcome Trust which was genotyped by the Wellcome Trust Case Control Consortium and the Type-1 Diabetes Genetics Consortium, sponsored by the National Institute of Diabetes and Digestive and Kidney Diseases, National Institute of Allergy and Infectious Diseases, National Human Genome Research Institute, National Institute of Child Health and Human Development and Juvenile Diabetes Research Foundation International.

#### (3) European Alzheimer’s Disease Initiative (EADI)

All AD cases were ascertained by neurologists from Bordeaux, Dijon, Lille, Montpelier, Paris, and Rouen, with clinical diagnosis of probable AD established according to the DSM-III-R and National Institute of Neurological and Communication Disorders and Stroke-Alzheimer’s Disease and Related Disorders Association (NINCDS-ADRDA) criteria^51,52^. Controls were recruited from Lille, Rouen, Nantes and from the 3C Study^51^. This cohort is a population-based, prospective study of the relationship between vascular factors and dementia. It has been carried out in three French cities: Bordeaux (southwest France), Montpelier (southeast France) and Dijon (central eastern France). A sample of non-institutionalized, subjects over 65 years was randomly selected from the electoral rolls of each city. Between January 1999 and March 2001, 9,686 subjects meeting the inclusion criteria agreed to participate. Following recruitment, 392 subjects withdrew from the study. Thus, 9,294 subjects were finally included in the study (2,104 in Bordeaux, 4,931 in Dijon and 2,259 in Montpellier). At 8 years of follow up, 664 individuals suffered from AD with 167 prevalent and 497 incident cases. The other individuals were considered as controls. 9863 DNA samples that passed DNA quality control were genotyped with Illumina Human 610-Quad BeadChips. Following quality control procedures, a final sample size of 5,803 3C individuals (387 AD cases and 5,416 controls, cohort dataset) and 2,298 non-3C individuals (1,420 AD cases and 878 controls, case-control dataset) was included in this study.

#### (4) Cohorts for Heart and Aging Research in Genomic Epidemiology (CHARGE)

Cardiovascular Health Study (CHS): The CHS is a prospective population-based cohort study of risk factors for vascular and metabolic disease that in 1989-90, enrolled adults aged ≥65 years, at four field centers located in North Carolina, California, Maryland and Pennsylvania. The original predominantly Caucasian cohort of 5,201 persons was recruited from a random sample of people on Medicare eligibility lists and an additional 687 African-Americans were enrolled subsequently for a total sample of 5,8882. DNA was extracted from blood samples drawn on all persons who consented to genetic testing at their baseline examination in 1989-90. In 2007-2008, genotyping was performed at the General Clinical Research Center’s Phenotyping/Genotyping Laboratory at Cedars-Sinai using the Illumina 370CNV Duo ® BeadChip system on 3,980 CHS participants who were free of cardiovascular disease (CVD) at baseline. The 1,908 persons excluded for prevalent CVD had prevalent coronary heart disease (n=1,195), congestive heart failure (n=86), peripheral vascular disease (n=93), valvular heart disease (n=20), stroke (n=166) or transient ischemic attack (n=56). Some persons had more than one reason to be excluded and for these individuals only the initial exclusionary event is listed. Because the other cohorts were predominantly white, the African American CHS participants were excluded from this analysis to limit errors secondary to population stratification. Among white participants genotyping was attempted in 3,397 participants and was successful in 3295 persons. After excluding persons that had either died prior to the start of the CHS cognition study in 1992 (see section 3 for details), could not be evaluated completely for baseline cognitive status, and persons that had dementia other than AD, a sample of 2,049 persons was available. The CHS study protocols were approved by the Institutional review boards at the individual participating centers.

The AD sample for this study included all prevalent cases identified in 1992 and incident events identified between 1992 and December 20063. Briefly, persons were examined annually from enrollment to 1999. The examination included a 30 minute screening cognitive battery. In 1992-94 and again, in 1997-99, participants were invited to undergo brain MRI and detailed cognitive and neurological assessment as part of the CHS Cognition Study. Persons with prevalent dementia were identified, and all others were followed until 1999 for the development of incident dementia and AD. Since then, CHS participants at the Maryland and Pennsylvania centers have remained under ongoing dementia surveillance^53^.

Beginning in 1988/89, all participants completed the Modified Mini-Mental State Examination (3MSE) and the DSST at their annual visits, and the Benton Visual Retention Test (BVRT) from 1994 to 1998. The Telephone Interview for Cognitive Status (TICS) was used when participants did not come to the clinic. Further information on cognition was obtained from proxies using the Informant Questionnaire for Cognitive Decline in the Elderly (IQCODE), and the dementia questionnaire (DQ). Symptoms of depression were measured with the modified version of the Center for Epidemiology Studies Depression Scale (CES-D). In 1991-94, 3,608 participants had an MRI of the brain and this was repeated in 1997-98. The CHS staff also obtained information from participants and next-of-kin regarding vision and hearing, the circumstances of the illness, history of dementia, functional status, pharmaceutical drug use, and alcohol consumption. Data on instrumental activities of daily living (IADL), and activities of daily living (ADL) were also collected.

Persons suspected to have cognitive impairment based on the screening tests listed above underwent a neuropsychological and a neurological evaluation. The neuropsychological battery included the following tests: the American version of the National Reading test (AMNART), Raven’s Coloured Progressive Matrices, California Verbal Learning Test (CVLT), a modified Rey-Osterreith figure, the Boston Naming test, the Verbal fluency test, the Block design test, the Trails A and B tests, the Baddeley & Papagno Divided Attention Task, the Stroop, Digit Span and Grooved Pegboard Tests. The results of the neuropsychological battery were classified as normal or abnormal (>1.5 standard deviations below individuals of comparable age and education) based on normative data collected from a sample of 250 unimpaired subjects. The neurological exam included a brief mental status examination, as well as a complete examination of other systems. The examiner also completed the Unified Parkinson’s Disease Rating Scale (UPDRS) and the Hachinski Ischemic Scale. After completing the neurological exam, the neurologist classified the participant as normal, having mild cognitive impairment (MCI), or dementia.

International diagnostic guidelines, including the NINCDS-ADRDA criteria for probable and possible AD and the ADDTC’s State of California criteria for probable and possible vascular dementia (VaD) with or without AD, were followed. CHS identified 3 subtypes: possible/probable AD without VaD (categorized as pure AD, included in all AD) and mixed AD (for cases that met criteria for both AD and VaD, included in all-AD), and, possible/probable VaD without AD (excluded from current study).

Framingham Heart Study (FHS): The FHS is a three-generation, single-site, community-based, ongoing cohort study that was initiated in 1948. It now comprises three generations of participants including the Original cohort followed since 1948 (n=5,209)^54^, their Offspring and spouses of the offspring (n=5,216) followed since 1971^55^; and children from the largest Offspring families enrolled in 2000 (Gen 3)^56^. Participants in the Original and Offspring cohorts are used in these analyses, but Gen 3 participants were not included since they are young (mean age 40±9 years in 2000) and none had developed Alzheimer’s Disease (AD). The Original cohort enrolled 5,209 men and women who comprised two-thirds of the adult population then residing in Framingham, Massachusetts. Survivors continue to receive biennial examinations. The Offspring cohort comprises 5,124 persons (including 3,514 biological offspring) who have been examined approximately once every 4 years. Almost all the FHS Original and Offspring participants are white/Caucasian. FHS participants had DNA extracted and provided consent for genotyping in the 1990s. All available eligible participants were genotyped at Affymetrix (Santa Clara, CA) through an NHLBI funded SNP-Health Association Resource (SHARe) project using the Affymetrix GeneChip ® Human Mapping 500K Array Set and 50K Human Gene Focused Panel ®. In 272 persons, small amounts of DNA were extracted from stored whole blood and required whole genome amplification prior to genotyping. Cell lines were available for most of the remaining participants. Genotyping was attempted in 5,293 Original and Offspring cohort participants, and 4,425 persons met QC criteria. Failures (call rate <97%, extreme heterozygosity or high Mendelian error rate) were largely restricted to persons with whole-genome amplified DNA and DNA extracted from stored serum samples. In addition, since the persons with whole genome amplified DNA represent a group of survivors who may differ from the others we included whole genome amplified status as a covariate in FHS analyses. After exclusion of prevalent dementia, dementia other than AD, and missing values, a sample of 2,208 participants was available for this project. The FHS component of this study was approved by the Institutional Review Board of the Boston Medical Center.

The Original cohort of the FHS has been evaluated biennially since 1948, was screened for prevalent dementia and AD in 1974-76 and has been under surveillance for incident dementia and AD since then^57–59^. The Offspring have been examined once every 4 years and have been screened for prevalent dementia with a neuropsychological battery and brain MRI^60,61^. In order to be consistent with the sampling frame for the AGES and CHS samples, we excluded FHS subjects with a baseline age <65 yrs at the time of DNA draw which was in the 1990s. To minimize survival biases, Original cohort and Offspring participants who developed dementia prior to the date of DNA draw were treated as prevalent cases, and subsequent events in the Original cohort occurring prior to December 2006 were included in the incident analyses.

At each clinic exam, participants receive questionnaires, physical examinations and laboratory testing; between examinations they remain under surveillance (regardless of whether or not they live in the vicinity) via physician referrals, record linkage and annual telephone health history updates. Methods used for dementia screening and follow-up have been previously described^57,62^. Briefly, surviving cohort members who attended biennial examination cycles 14 and 15 (May 1975-November 1979) were administered a standardized neuropsychological test battery to establish a dementia-free cohort. Beginning at examination cycle 17 (1982), the MMSE was administered biennially to the cohort. A MMSE score below the education-specific cutoff score, a decline of 3 or more points on subsequent administrations, a decline of more than 5 points compared with any previous examination, or a physician or family referral prompted further in-depth testing. The Offspring cohort that was enrolled in 1971 has undergone 8 re-examinations, one approximately every 4 years. Starting at the 2nd Offspring examination, participants were questioned regarding any subjective memory complaints and since the 5th Offspring examination participants have been administered the MMSE at each visit. In addition concurrent with the 7th and 8th Offspring examinations (between 1999 and 2004 and then again between 2005 and 2009) surviving Original cohort and all eligible and consenting Offspring participants have undergone volumetric brain MRI and neuropsychological testing^60,61^. The neuropsychological test battery included the Reading subtest of the Wide Range Achievement Test (WRAT-3), the Logical Memory and the Paired Associates Learning tests from the Wechsler Memory Scale, the Visual Reproduction and Hooper Visual Organization Tests, Trails A and B, the Similarities subtest from the Wechsler Adult Intelligence test, the 30-iterm version of the Boston Naming Test and at the second assessment only, the Digit Span, Controlled Word Association and Clock Drawing Tests. Offspring participants suspected to have cognitive impairment based on their MMSE scores, participant, family or physician referral, hospital records or performance in the neuropsychological test battery described above were referred for more detailed neuropsychological and neurological evaluation.

Each participant thus identified underwent baseline neurologic and neuropsychological examinations. Neurologists (trained in geriatric behavioral assessment) supplemented their clinical assessment with a few structured cognitive tests and administered the Clinical Dementia Rating (CDR). Persons were reassessed systematically for the onset of at least mild dementia. A panel consisting of at least 1 neurologist (S.A., P.A.W., or S.S.) and 1 neuropsychologist (R.A.) reviewed all available medical records to arrive at a final determination regarding the presence or absence of dementia, the date of onset of dementia, and the type of dementia. For this determination, we used data from the neurologist’s examination, neuropsychological test performance, Framingham Study records, hospital records, information from primary care physicians, structured family interviews, computed tomography and magnetic resonance imaging records, and autopsy confirmation when available. All individuals identified as having dementia satisfied the DSM-IV criteria, had dementia severity equivalent to a CDR of 1 or greater, and had symptoms of dementia for at least 6 months. All individuals identified as having Alzheimer-related dementia met the NINCDS-ADRDA criteria for definite, probable, or possible AD. Vascular Dementia was diagnosed using the ADDTC criteria but the presence of vascular dementia did not disqualify a participant from obtaining a concomitant diagnosis of AD if indicated. The recruitment of Original cohort participants at FHS had occurred long before the DNA collection with the result that the majority of dementia events in the FHS (although ascertained prospectively) were prevalent at the time of DNA collection or these persons had died prior to DNA draw and were thus excluded from analyses of incident disease. Due to the limited number of incident dementia and AD events in the Framingham Offspring only the Original cohort were included in our analyses of incident events.

Rotterdam Study: The Rotterdam Study enrolled inhabitants from a district of Rotterdam (Ommoord) aged ≥55 years (N=7,983, virtually all white) at the baseline examination in 1990-93 when blood was drawn for genotyping^63^. It aims to examine the determinants of disease and health in the elderly with a focus on neurogeriatric, cardiovascular, bone, and eye disease. All inhabitants of Ommoord aged ≥55 years (n=10,275) were invited and the participation rate was 78%. All participants gave written informed consent to retrieve information from treating physicians. Baseline measurements were obtained from 1990 to 1993 and consisted of an interview at home and two visits to the research center for physical examination. Survivors have been re-examined three times: in 1993-1995, 1997-1999, and 2002-2004. All persons attending the baseline examination in 1990-93 consented to genotyping and had DNA extracted. This DNA was genotyped using the Illumina Infinium II HumanHap550chip v3·0 ® array in 2007-2008 according to the manufacturer’s protocols. Genotyping was attempted in persons with high-quality extracted DNA (n=6,449). From these 6,449, samples with low call rate (<97.5%, n=209), with excess autosomal heterozygosity (>0.336, n=21), with sex-mismatch (n=36), or if there were outliers identified by the IBS clustering analysis (>3 standard deviations from population mean, n=102 or IBS probabilities >97%, n=129) were excluded from the study population with some persons meeting more than one exclusion criterion; in total, 5,974 samples were available with good quality genotyping data, 42 persons were excluded since they did not undergo cognitive screening at baseline, hence their cognitive status was uncertain. An additional 61 persons were excluded because they suffered from dementia other than AD at baseline. After exclusion of prevalent dementia, a sample of 5752 persons was available. The Rotterdam Study (including its brain magnetic resonance imaging (MRI) and neurological components) has been approved by the institutional review board (Medical Ethics Committee) of the Erasmus Medical Center and the Netherlands Ministry of Health, Welfare and Sports Participants were screened for prevalent dementia in 1990-93 using a three-stage process; those free of dementia remained under surveillance for incident dementia, a determination made using records linkage and assessment at three subsequent re-examinations. We included all prevalent cases and all incident events up to 31st December 2007.

Screening was done with the MMSE and GMS organic level for all persons. Screen-positives (MMSE <26 or Geriatric Mental Schedule (GMS) organic level >0) underwent the CAMDEX. Persons who were suspected of having dementia underwent more extensive neuropsychological testing. When available, imaging data were used. In addition, all participants have been continuously monitored for major events (including dementia) through automated linkage of the study database with digitized medical records from general practitioners, the Regional Institute for Outpatient Mental Health Care and the municipality. In addition physician files from nursing homes and general practitioner records of participants who moved out of the Ommoord district were reviewed twice a year. For suspected dementia events, additional information (including neuroimaging) was obtained from hospital records and research physicians discussed available information with a neurologist experienced in dementia diagnosis and research to verify all diagnoses. Dementia was diagnosed in accordance with internationally accepted criteria for dementia (Diagnostic and Statistical Manual of Mental Disorders, Revised Third Edition, DSM-III-R), and AD using the NINCDS-ADRDA criteria for possible, probable and definite AD. The National Institute of Neurological Disorders and Stroke–Association Internationale pour la Recherche et l’Enseignement en Neurosciences (NINDS-AIREN) criteria were used to diagnose vascular dementia. The final diagnosis was determined by a panel of a neurologist, neurophysiologist, and research physician and the diagnoses of AD and VaD were not mutually exclusive.

#### (5) Power Calculation

To determine the power to detect genetic variants associated with age at onset, we ran analyses using Proc Power in SAS. The analysis was run using minor allele frequencies ranging from 0.05 to 0.50, OR 1.1 to 1.75 and sample size of 45,000. Other factors, such as genetic heterogeneity and gene-environment interaction are likely to affect these estimates. Alpha was adjusted to 5×10^−8^. For variants with a MAF of 0.15, we would have approximately 80% power to detect effects for OR > 1.23 or < 0.81; for variants with a MAF of 0.3, we would have approximately 80% power to detect effects for OR > 1.18 (or < 0.85).

### CSF biomarker datasets

CSF samples were obtained from the Knight-ADRC (N=805), ADNI-1 (N=390), ADNI-2 (N=397), the Biomarkers for Older Controls at Risk for Dementia (BIOCARD) (N=184), Mayo Clinic (N=433), Lund University (Swedish) (N=293), University of Pennsylvania (Penn) (N=164), University of Washington (N=375), The Parkinson’s Progression Markers Initiative (500) and Saarland University (German) (N=105).

Cases were diagnosed with dementia of the Alzheimer’s type (DAT) according to the NINCDS-ADRDA^20^. Control individuals were evaluated using the same criteria and showed no symptoms of cognitive impairment. All participants provided written informed consent and the ethics committee approved the informed consent procedure (IRB ID #: 201105364). 787 additional samples with biomarker data used in the analyses were obtained from the ADNI database (adni.loni.usc.edu). ADNI was launched in 2003 as a public-private partnership, led by Principal Investigator Michael W. Weiner, MD. The primary goal of ADNI has been to test whether serial magnetic resonance imaging (MRI), positron emission tomography (PET), other biological markers, and clinical and neuropsychological assessment can be combined to measure the progression of mild cognitive impairment (MCI) and early Alzheimer’s disease (AD).

CSF in all studies was collected in a standardized manner^12,64–67^. Biomarker measurements within each study were conducted using internal standards and controls to achieve consistency and reliability. However, differences in the measured values between studies were observed which are likely due to differences in the antibodies and technologies used for quantification (standard ELISA with Innotest for Knight-ADRC, UW, Swedish, German, and Mayo versus Luminex with AlzBio3 for ADNI-1, ADNI-2, BIOCARD and Penn), ascertainment and/or handling of the CSF after collection. CSF Aβ_42_ and ptau_181_ values were log transformed in order to approximate a normal distribution. Because the CSF biomarker values were measured using two different platforms (standard ELISA with Innotest and Luminex with AlzBio3), we did not combine the raw data. For the combined analyses, we standardized the mean of the log-transformed values from each dataset to zero. No significant differences in the transformed and standardized CSF values were found between cohorts. We also performed meta-analyses for the most significant SNPs by combining the P values for each independent dataset using METAL^68^. No major differences were found between the joint-analyses and the meta-analyses.

### Quality Control

For survival analysis, we excluded cases with AAO below 60 and cases with prevalent stroke. For CSF analysis, individuals under age 45 years were removed because prior studies have demonstrated that the relationship between CSF Aβ_42_ levels and age appears to differ in individuals below 45 years vs. those above 45 years^69^. Of the remaining individuals in both analyses, we excluded individuals who had > 5% missing genotype rates, who showed a discrepancy between reported sex and sex estimated on the basis of genetic data, or who showed evidence of non-European ancestry based on principal component analysis using PLINK1.9^70^.

We identified unanticipated duplicates and cryptic relatedness using pair-wise genome-wide estimates of proportion identity by descent (IBD) using PLINK. When duplicate samples or a pair of samples with cryptic relatedness was identified, the sample with the lower genotyping call rate was removed. We excluded potentially related individuals so that all remaining individuals have kinship coefficient below 0.05. Finally, we excluded individuals with missing disease status, age or gender information.

To control for genotype quality, we excluded SNPs with missing genotypes in > 5% of individuals in each dataset for survival analysis, and > 2% for CSF association analysis. For the EADI cohort, variants with minor allele frequency < 1%, Hardy-Weinberg P value < 1×10^−6^ and missingness > 2% were removed prior to imputation. Genome-wide genotype imputation was performed using IMPUTE2^71^ with 1000 Genomes reference haplotypes. We excluded imputed SNPs with an IMPUTE2 quality score < 0.5 for survival analysis. For CSF association, we excluded SNPs with an IMPUTE2 quality score of < 0.3 since the dataset was only used for follow-up. In the ADGC, GERAD, CHARGE, and CSF datasets, we then removed SNPs that failed the Hardy-Weinberg equilibrium in controls calculated based on the imputed best-guess genotypes using a P value threshold of 1×10^−6^. We excluded SNPs with minor allele frequency ≤ 0.02. Finally, we excluded SNPs with available statistics in only one consortium dataset in the meta-analysis. The number of filtered samples and SNPs in each of the above steps are recorded in **Supplementary Table 1**.

### Genome-wide survival association study

We conducted a genome-wide Cox proportional hazards regression^72^ assuming an additive effect from SNP dosage. The Cox proportional hazard regression was implemented in the R survival analysis package. We incorporated sex, site and the first three principal components from EIGENSTRAT^30^ in all our regression models to control for their effects. For EADI, sex and four principal components were included in the model. For the Cox model, the time scale is defined as age in years, where age is age at onset for cases and age at last assessment for controls. The formula applied is as followed:

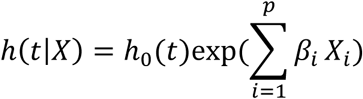

where X=(X1, X2, …, Xp) are the observed values of covariates for subject i. The Cox model has previously been shown to be applicable to case-control datasets without an elevated type 1 error rate nor overestimation in effect sizes^73,74^. The model assumes log-linearity and proportional hazards. The assumption of log-linearity is common in the additive logistic regression used in a typical GWAS. We validated the assumption of proportional hazards assumed by the Cox model by conducting the Schoenfeld test in the 22 prioritized SNPs. None of the SNPs has a Schoenfeld P value, which is the P value for Pearson product-moment correlation between the scaled Schoenfeld residuals and time, lower than 0.035 (multiple test correction threshold=0.00227) in any of the 7 cohorts. Further, only 3 out of the 148 P values were less than 0.05, suggesting that the time proportionality assumption is unlikely to be violated in these associations (**Supplementary Table 1**). Similarly, the Schoenfeld test was conducted for all 22 SNP association models on the covariates in the ADGC and GERAD cohort (**Supplementary Table 1**). We also examined the effect sizes of our candidate SNPs in these cohorts and found consistent effect sizes (Supplementary Fig. 3) in the 3 retrospective case-control cohorts (ADGC, GERAD, EADI case-control) and 4 prospective cohorts (EADI-prospective, CHARGE FHS, CHS and Rotterdam).

After the analysis of each dataset, we carried out an inverse-variance meta-analysis on the results using METAL^26^, applying a genomic control to adjust for inflation in each dataset. Of the 751 suggestive SNPs (P <1×10^−5^), we found these SNPs to show lower standard errors and confidence intervals with the increasing number of cohorts showing consistent directionalities of effect. Particularly, the average standard error for SNPs showing 1 to 7 consistent directionalities ranges from 0.171, 0.109, 0.0744, 0.0346, 0.0234, 0.0173 to 0.01795 (Supplementary Fig. 1b). Thus, we limited our final analysis to SNPs that showed consistent directionalities of effect in at least 6 out of the 7 datasets included in the meta-analysis. The association graphs of results from loci of interest were plotted using LocusZoom^75^.

### CSF biomarker association analysis

For the CSF datasets, we performed multivariate linear regression for CSF Aβ_42_ and tau, and ptau_181_ association adjusting for age, gender, site, and the first three principal components using PLINK.

### eQTL analysis

We examined the effect of top survival and CSF SNPs on gene expression using published databases. For general brain expression eQTL analysis, we queried the BRAINEAC eQTL data provided by the UK human Brain Expression Consortium (see URLs).

We conducted leukocyte-specific analysis using the Cardiogenics dataset^27^ composed of 738 monocytes and 593 macrophages samples. For each probeset – imputed SNP pair, a simple linear regression was used to analyze the data separately for monocytes and macrophages:

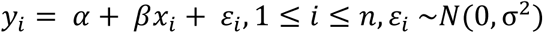

where i is the subject index, x is the effective allele copy number, and y_i_ is the covariates-adjusted, inverse-normal transformed gene expression. Significance of *cis* (SNP within ± 1Mb of the closest transcript end) eQTL effects were quantified with a Wald test on the ordinary Least Squares (OLS) estimator of the coefficient β, obtained with R. The distribution of the Wald test P values under the null hypothesis of no correlation between genotype and gene expression was estimated by rerunning the same analysis on a null dataset obtained by permuting the expression samples identifiers. For additional monocyte eQTL analysis, we queried statistics from Fairfax et al.^28^ to validate findings in the Cardiogenics dataset.

For conditional analysis, we performed analysis for *SPI1* (probe: ILMN_1696463) against all SNPs within ± 2Mb from the closest transcript end, by including the following SNPs effective allele copy numbers as covariates in the linear regression model, one at a time: rs1057233, rs10838698, rs7928163, rs10838699, rs10838725, rs1377416. Significance was again assessed with a two-sided Wald test on the OLS estimator of the coefficient β.

### Gene expression analysis in human and mouse brain cell types

Cell-type specific gene expression in the human and mouse brain was queried from brain RNA-Seq databases described in Zhang et al.^29,30^ and Bennett et al.^31^ and plotted using custom R scripts (see URLs). The mouse astrocytes-FACS and astrocytes-immunopanned in mouse were collapsed into a single astrocyte cell type.

### Epigenetic analysis in human myeloid cell types

We utilized HaploReg^34^ to annotate the regulatory element of the significantly associated SNPs and their tagging SNPs. The myeloid chromatin marks/states and PU.1 ChIP-Seq data at genetic loci were further examined through the Washington University Epigenome browser^76^ using the public Roadmap Epigenomics Consortium public tracks hub as well as custom track hubs for human monocytes and macrophages (hg19) (see URLs).

### Colocalization (coloc and SMR/HEIDI) analyses

Colocalization analysis of genetic variants associated with AD and myeloid gene expression was performed using AAOS GWAS SNP and myeloid (monocyte and macrophage) eQTL datasets from Cardiogenics as inputs. Overlapping SNPs were retained within the hg19 region chr11:47100000-48100000 for the *SPI1/CELF1* locus, chr11:59500000-60500000 for the *MS4A* locus, and chr1:169300000-170300000 for the *SELL* locus. Colocalization analysis of AD-and gene expression-associated SNPs was performed using the ‘coloc.abf’ function in the ‘coloc’ R package (v2.3-1). Default settings were used as prior probability of association: 1×10^−4^ for trait 1 (gene expression), 1×10^−4^ for trait 2 (AD) and 1×10^−5^ for both traits. SMR/HEIDI (v0.65) analysis was performed as described in Zhu et al.^23^ and the companion website (see URLs). The ADGC subset of the IGAP GWAS dataset was used to perform the LD calculations.

### Partitioned heritability analysis using LD score regression

We used LDSC (LD SCore, v1.0.0) regression analysis^25^ to estimate heritability of AD and schizophrenia from GWAS summary statistics (excluding the APOE [chr19:45000000-45800000] and MHC/HLA [chr6:28477797-33448354] regions) partitioned by PU.1 ChIP-Seq binding sites in myeloid cells, as described in the companion website (see URLs) and controlling for the 53 functional annotation categories of the full baseline model. GWAS summary statistics for AD and schizophrenia (SCZ) were downloaded from the IGAP consortium^1^ (stage 1 dataset) and the Psychiatric Genomics Consortium (PGC)^26^ (pgc.cross.scz dataset), respectively (see URLs). SPI1 (PU.1) bindings sites were downloaded as filtered and merged ChIP-Seq peaks in BED format from the ReMap database^77^ (GEO:GSE31621, SPI1, blood monocyte and macrophage datasets^35^). SPI1 (PU.1) and POLR2AphosphoS5 binding sites were downloaded as broad ChIP-Seq peaks in BED format from the Encode portal^7837^ (DCC:ENCSR037HRJ; GEO:GSE30567; HL60 dataset) (see URLs).

### Phagocytosis assay

BV2 mouse microglial cell line was kindly provided by Marc Diamond (UT Southwestern Medical Center). BV2 cells were cultured in DMEM (Gibco 11965) supplemented with 5% FBS (Sigma F4135) and 100 U/ml penicillin-streptomycin (Gibco 15140). Routine testing of cell lines using MycoAlert PLUS mycoplasma detection kit (Lonza) showed that BV2 cells were negative for mycoplasma contamination. pcDNA3-FLAG-PU.1 was a gift from Christopher Vakoc^79^ (Addgene plasmid 66974). pGFP-V-RS with either non-targeting shRNA or PU.1-targeting shRNAs was purchased from OriGene Technologies (TG502008). The pHrodo red zymosan conjugate bioparticles from Thermo Fisher (P35364) were used to assess phagocytic activity. For transient transfections, 200,000 cells were seeded in a 24-well plate. On the next day, cells were washed with PBS (Gibco 14190) and medium was changed to 400 μl DMEM supplemented with 2% FBS without antibiotic. Transfection mixes of 0.5 μg pcDNA3 or 0.5 μg pcDNA3-FLAG-PU.1 with 0.5 μg pCMV-GFP for overexpression of mouse PU.1 and 1μg pGFP-V-RS-shSCR, -shA, -shB and -shD for knock-down of mouse PU.1 were prepared with 2 μl of Lipofectamine 2000, incubated for 20 min at room temperature and added to each well. After 8 hours of incubation 1 ml of growth medium was added to each well and plates were incubated for 2 days. Then the medium was replaced with 500 μl of fresh medium, and 25 μg of bioparticles were added to cells for 3 hour incubation. Bioparticles uptake was verified with a fluorescent microscope; then the cells were collected with trypsin (Gibco #25200), washed with PBS once and re-suspended in 500 μl PBS with 1% BSA. Cells were kept on ice and phagocytic activity was analyzed on an LSR II flow cytometer (BD Biosciences). At least 30,000 events were collected in each experiment, gated on FSC-A/SSC-A and further on FSC-A/FSC-W dot plot to analyze populations of viable single cells. Data were quantified using FCS Express 5 (De Novo Software) and GraphPad Prism 7 (GraphPad Software). Cells pretreated with 2 μM Cytochalasin D for 30 minutes before and during the uptake of bioparticles were used as a negative control. The population of GFP^+^/pHrodo^+^ cells in each condition was used to quantify the phagocytic index: percentage of pHrodo^+^ cells in GFP^+^ gated population x geometric mean pHrodo intensity / 10^6^; and represented as phagocytic activity. Three independent experiments were performed with two technical replicates without randomization of sample processing, n=3. Researcher was not blinded to the samples identification. Differences between the means of preselected groups were analyzed with one-way ANOVA and Sidak’s post hoc multiple comparisons test between selected groups, with a single pooled variance. Values of Cytochalasin D-treated cells were excluded from the statistical analysis. Adjusted P values for each comparison are reported, non-significant differences are not reported.

### Western blotting

BV2 cells transiently transfected as described for the phagocytosis assay were collected with trypsin after 48 hours of incubation, washed with PBS and re-suspended in PBS with 1% BSA. Cells from the same treatment were pooled and sorted on FACSARIA III (BD Biosciences) into GFP^+^ and GFP^−^ populations, pelleted at 2,000 rpm and lysed in RIPA buffer (50 mM Tris-HCl pH 7.4, 150 mM NaCl, 1% NP-40, 0.5% sodium deoxycholate, 0.1% SDS and Complete protease inhibitor tablets (Roche)) with one freeze-thaw cycle and 1 hour incubation on ice. Protein concentration was quantified using the BCA kit (Thermo Fisher #23225). Equal amounts of protein were separated by electrophoresis in Bolt 4–12% Bis-Tris Plus gels with MOPS SDS running buffer and transferred using the iBlot 2 nitrocellulose transfer stack. Membranes were blocked and probed with antibodies against PU.1 (Cell Signaling #2266) and β-Actin (Sigma #A5441) in 3% non-fat dry milk in TBS / 0.1% Tween-20 buffer. Secondary antibody staining was visualized using WesternBright ECL HRP Substrate Kit (Advansta K-12045) and ChemiDoc XRS+ (BioRad). Images were quantified using ImageJ (NIH) and GraphPad Prism 7 (GraphPad Software). Two independent experiments were performed without randomization of sample processing, n=2. Researcher was not blinded to the samples identification. Differences between every group mean were analyzed with one-way ANOVA and Sidak’s post hoc multiple variance test between selected groups, with a single pooled variance. Adjusted P values for each comparison are reported.

### Quantitative PCR

Sorted GFP^+^ BV2 cells after overexpression or knock-down of PU.1 were collected as described for western blotting. Cell pellets were lysed in QIAzol reagent and RNA was isolated with RNAeasy Mini kit according to the manufacturer’s instructions (Qiagen) including the Dnase treatment step with RNase-free DNase set (Qiagen). Quantities of RNA were measured using Nanodrop 8000 (Thermo Scientific) and reverse transcription was performed with 1-2 μg of total RNA using High-Capacity RNA-to-cDNA kit (Thermo Fisher Scientific). qPCR was performed on QuantStudio 7 Flex Real-Time PCR System (Thermo Fisher Scientific) using Power SYBR Green Master Mix (Applied Biosystems) with one-step PCR protocol. 3 ng of cDNA was used for all genes except *Ms4a4a* when 24 ng of cDNA was used in a 10 μl reaction volume. Primers were from PrimerBank^80^ or designed using Primer-BLAST program (NCBI) and are listed in **Supplementary Table 14**. Ct values were averaged from two technical replicates for each gene. Geometric mean of average Ct for the housekeeping genes *GAPDH*, *B2M* and *ACTB* was used as a reference that was subtracted from the average Ct for a gene of interest (dCt). Gene expression levels were log transformed (2^-dCt^) and related to the combined mean values of pcDNA3 and pGFP-V-RS-shSCR control samples in each sort giving relative expression for each gene of interest. Data were visualized in GraphPad Prism 7 (GraphPad Software). Four independent experiments were performed without randomization of sample processing, n=4. Researcher was not blinded to the sample identity. Differences between means were analyzed using one-way ANOVA and Dunnett’s post hoc multiple comparisons test between experimental and control groups, with a single pooled variance. Adjusted P values for each comparison are reported in **Supplementary Table 13**.

### Data availability

Summary statistics for the genome-wide survival analyses are posted on the NIA Genetics of Alzheimer’s Disease Data Storage (NIAGADS, see URLs).

### Code availability

Codes for analyses are available at a public GitHub repository (https://github.com/kuanlinhuang/AD_SPI1_project).

## URLs

BRAINEAC, http://caprica.genetics.kcl.ac.uk/BRAINEAC; LDSC software, http://www.github.com/bulik/ldsc; baseline and cell type group annotations, http://data.broadinstitute.org/alkesgroup/LDSCORE/; stratified LD score regression companion website, https://github.com/bulik/ldsc/wiki/Partitioned-Heritability; SMR/HEIDI software and companion website, http://cnsgenomics.com/software/smr; Brain RNA-Seq, http://web.stanford.edu/group/barres_lab/brainseq2/brainseq2.html; WashU EpiGenome Browser, http://epigenomegateway.wustl.edu/browser; custom tracks for human monocytes and macrophages, http://www.ag-rehli.de/TrackHubs/hub_MOMAC.txt; International Genomics of Alzheimer’s Project (IGAP) http://web.pasteur-lille.fr/en/recherche/u744/igap/igap_download.php; Psychiatric Genomics Consortium (PGC) https://www.med.unc.edu/pgc/results-and-downloads; ReMap database http://tagc.univ-mrs.fr/remap; Encode portal https://www.encodeproject.org/; NIAGADS, https://www.niagads.org.

---

**Supplementary Figure 1.**
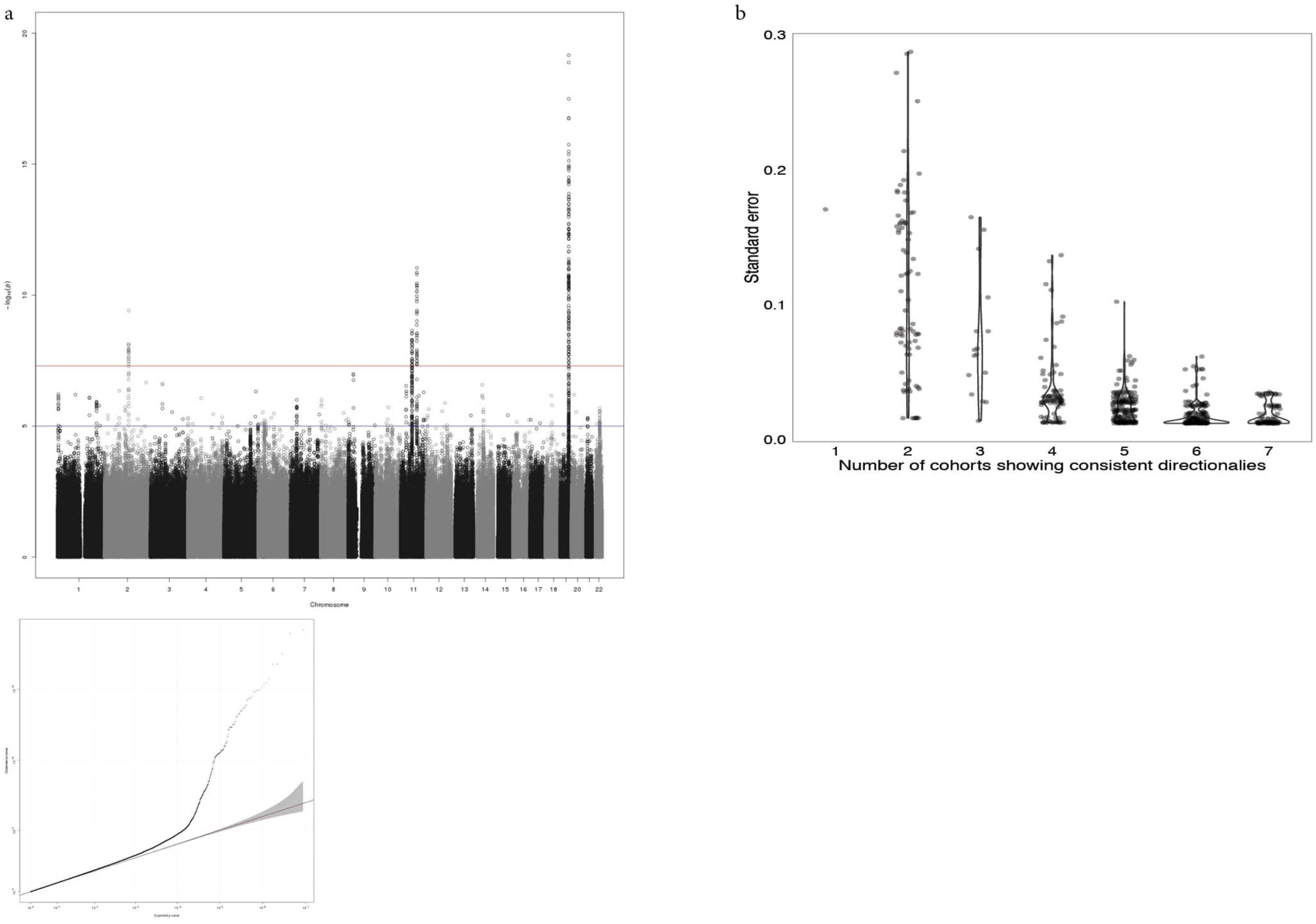
Result and quality control analysis of the IGAP AD-survival meta-analysis. (a) Manhattan plot and QQ-plot of the GWAS. The final meta-analysis showed little evidence of genomic inflation (λ=1.026). (b) The average standard error versus the number of cohorts with consistent directionalities of effect sizes.

**Supplementary Figure 2.**
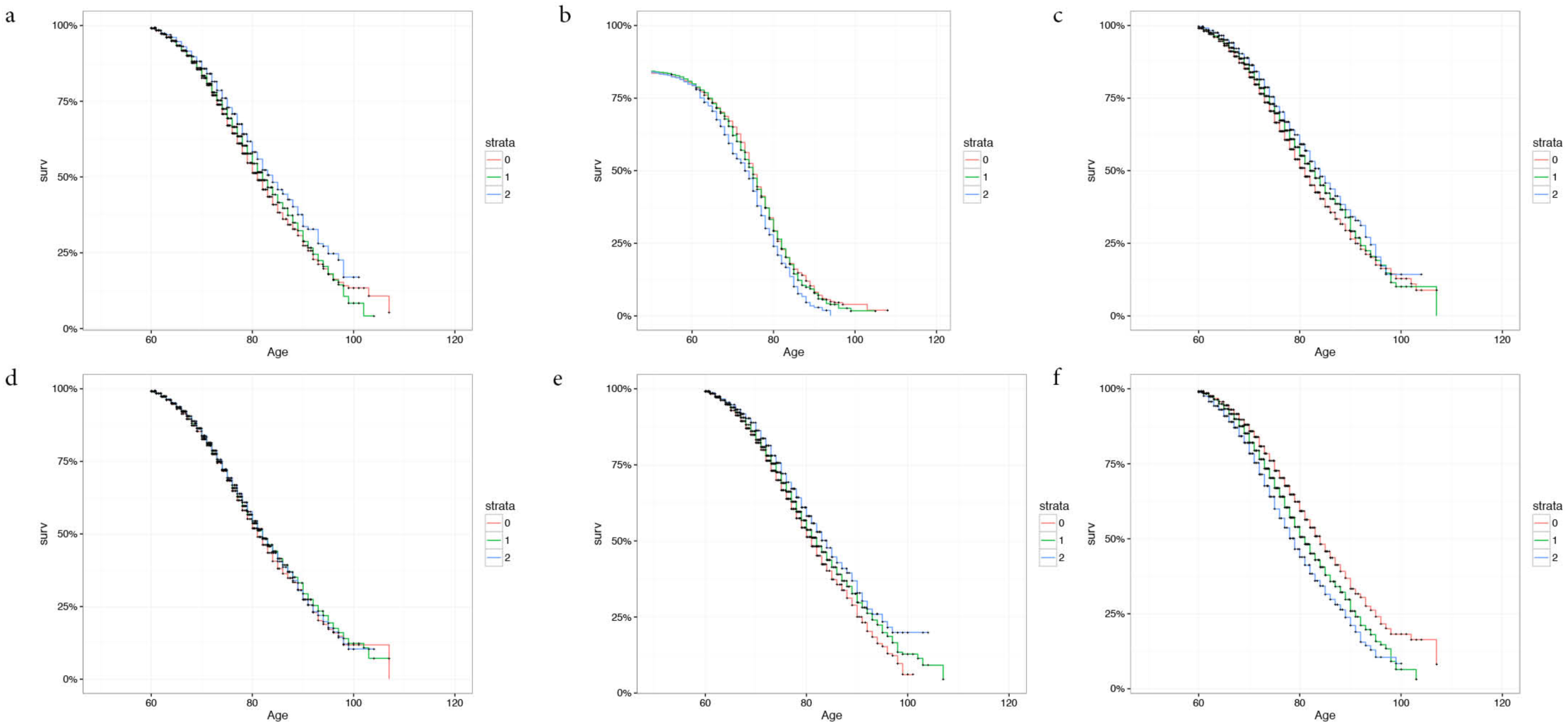
Kaplan-Meier plots of AAOS associations. Kaplan-Meier plots of survival analysis associations in the ADGC cohort of (a) rs1057233, (b) rs10919252, (c) rs567075, (d) rs7867518, (e) rs7930318, (f) rs4803758.

**Supplementary Figure 3.**
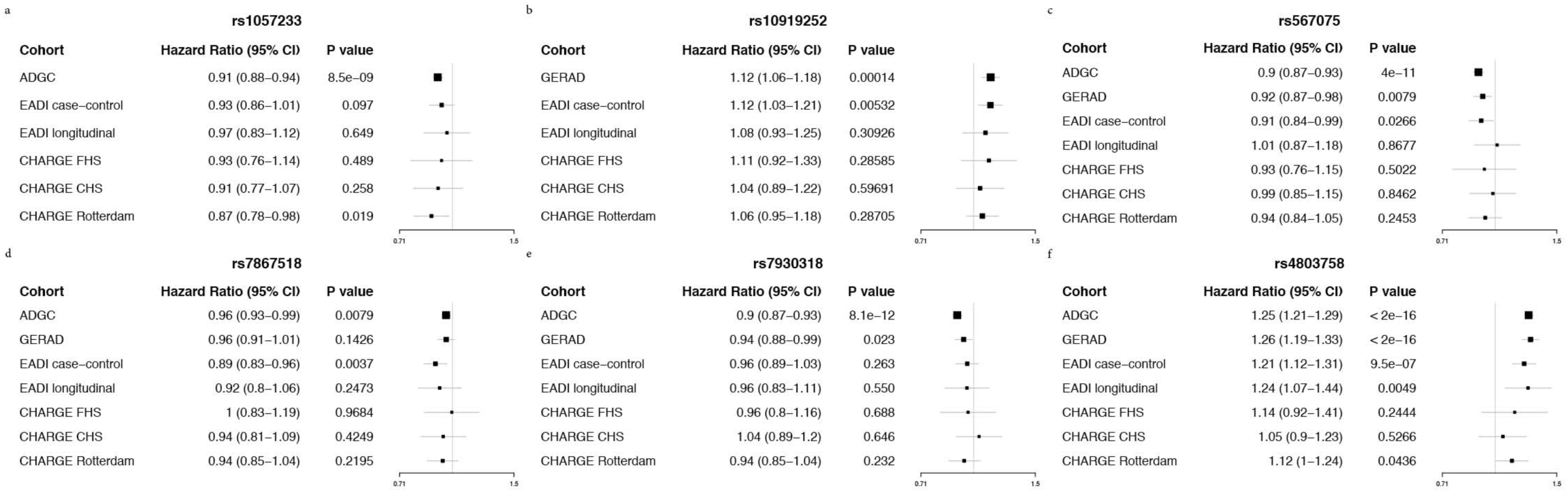
Forest plots of AAOS associations. Forest plots of survival analysis associations across IGAP cohorts of (a) rs1057233, (b) rs10919252, (c) rs567075, (d) rs7867518, (e) rs7930318, (f) rs4803758.

**Supplementary Figure 4.**
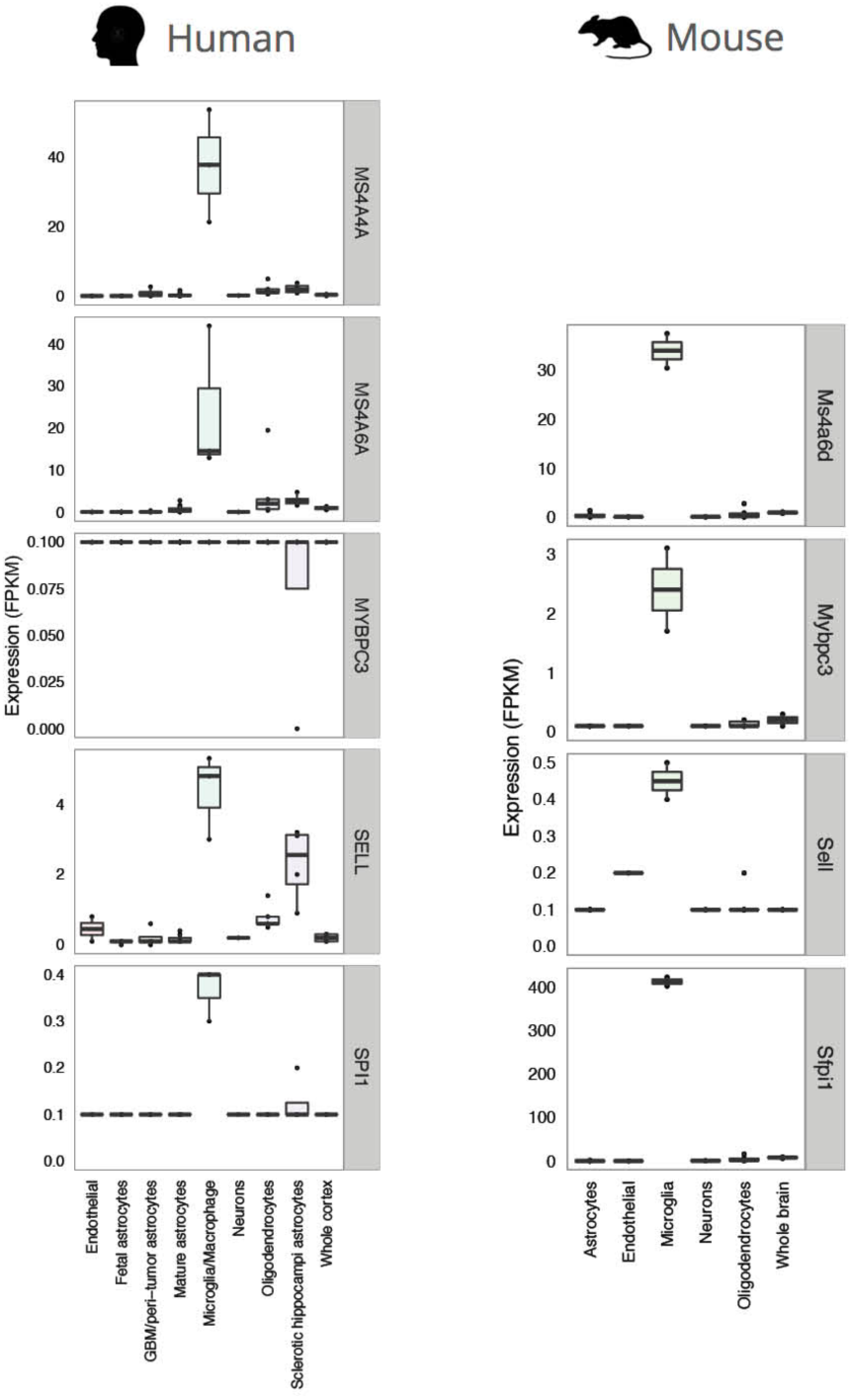
Cell-type specific expression of eQTL-associated genes in brain. Cell-type specific expression of *MS4A4A* (no mouse homolog available), *SPI1, MYBPC3, MS4A6A* and *SELL* in human and mouse brains based on the brain RNA-Seq database.

**Supplementary Figure 5.**
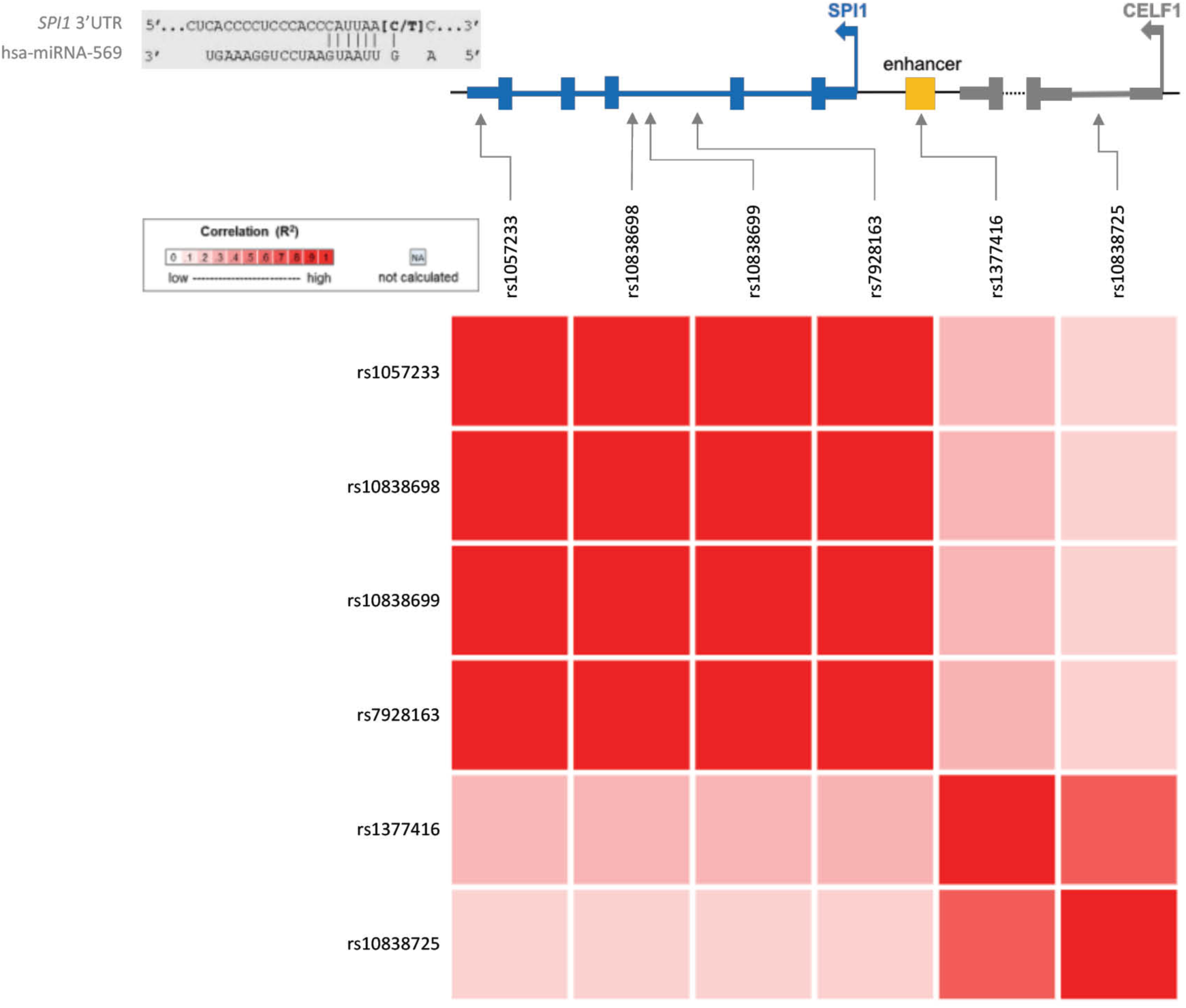
Linkage disequilibrium (LD) plot of SNPs of interest in the *SPI1/CELF1* locus.

**Supplementary Figure 6.**
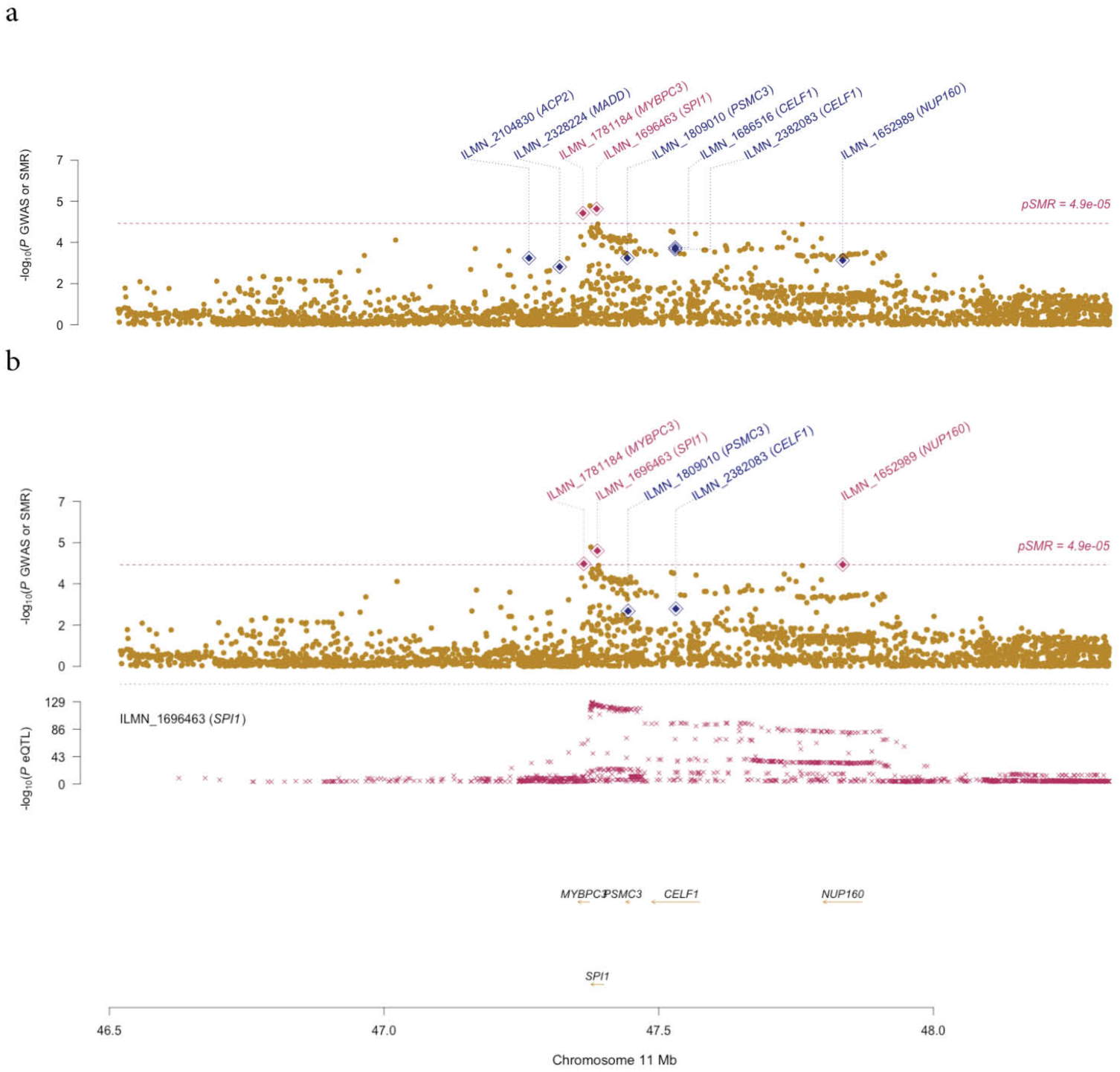
SMR plots showing the associations at the *SPI1/CELF1* locus. SMR plots showing the associations at the *SPI1/CELF1* locus from AAOS GWAS and eQTLs in (a) monocytes and (b) macrophages.

**Supplementary Figure 7.**
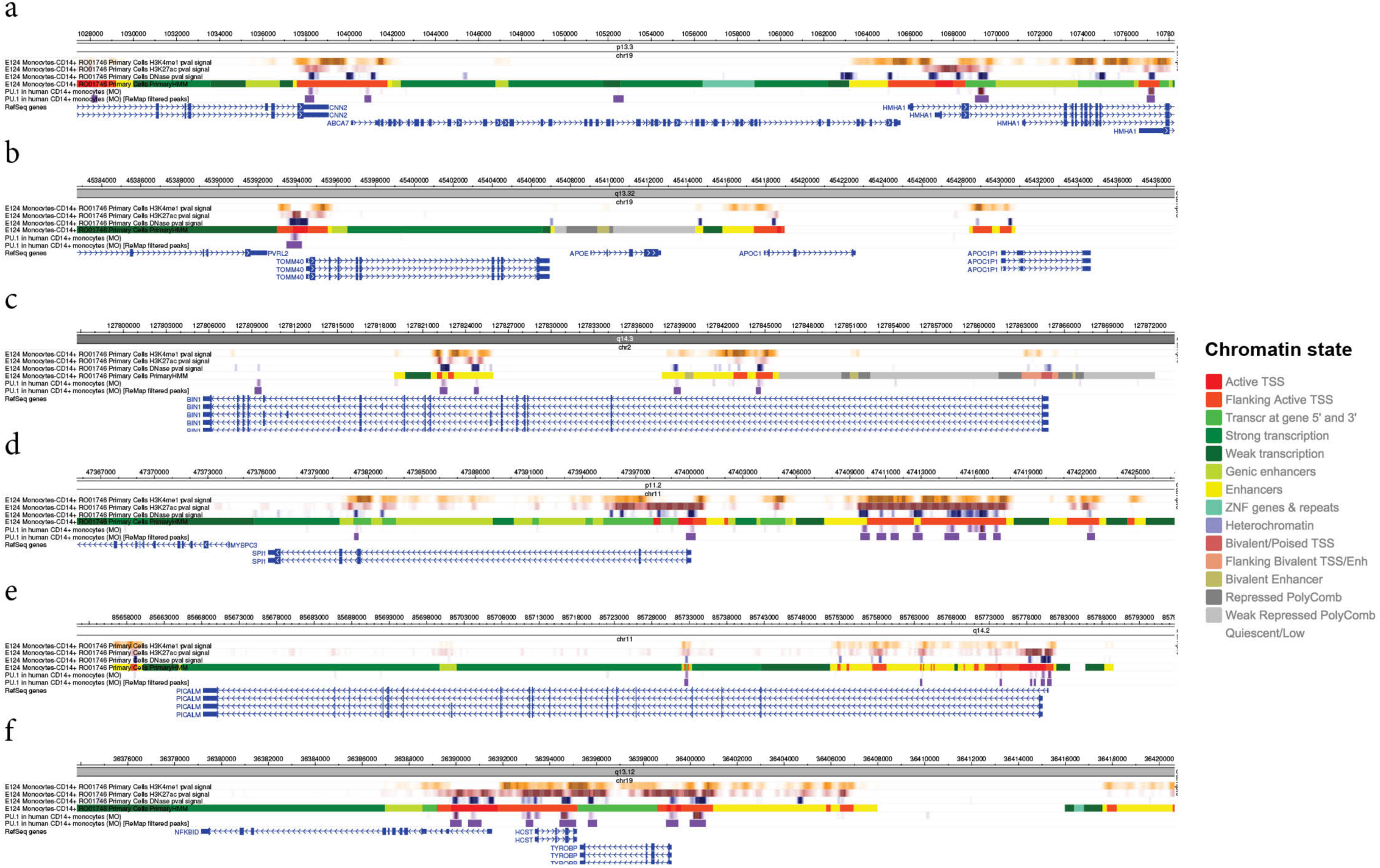
*SPI1* (PU.1) ChIP-Seq binding sites and other epigenetic signatures at AD-associated loci in human CD14+ monocytes. PU.1 binding sites, DNase I hypersensitive sites, histone modifications, and chromatin states at the locus of **(a)** *ABCA7*, **(b)** *APOE*, **(c)** *BIN1*, **(d)** *SPI1*, **(e)** *PICALM*, and **(f)** *TYROBP*.

**Supplementary Figure 8.**
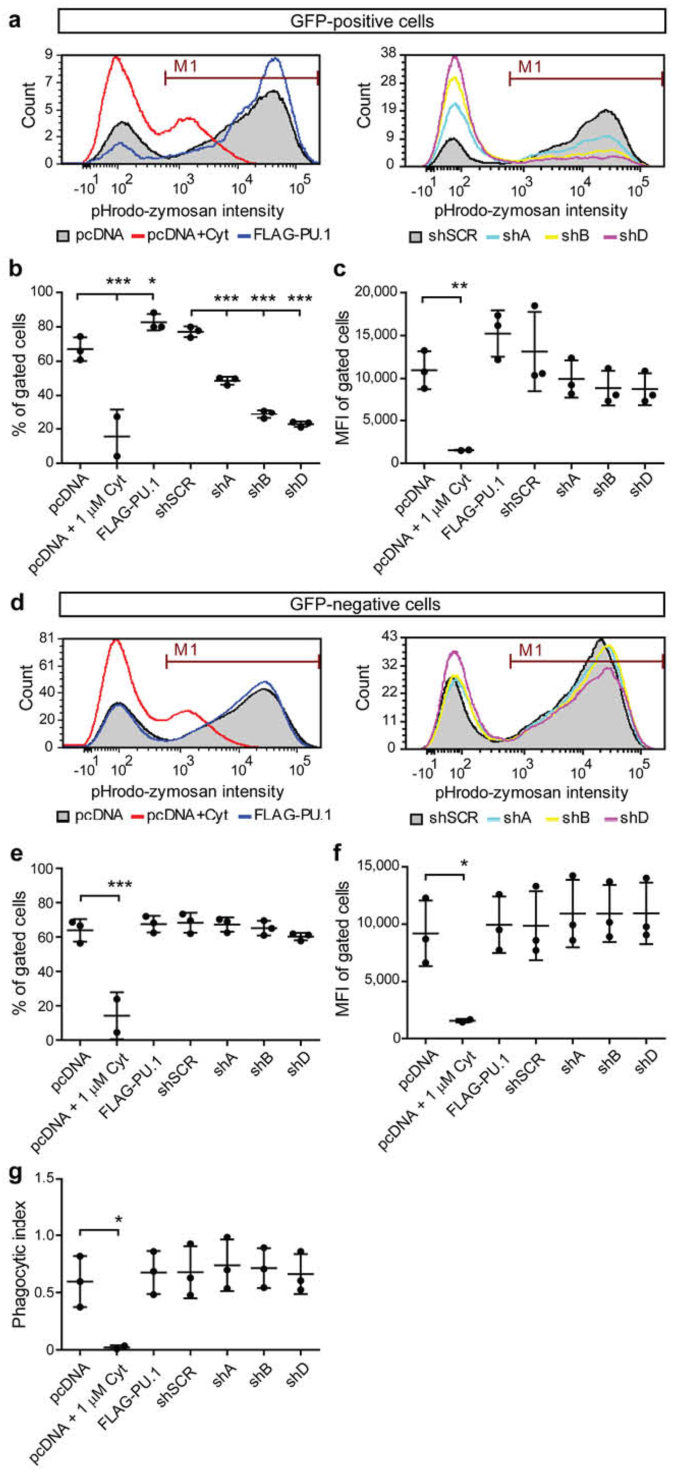
Analysis of phagocytosis in BV2 microglial cells. **(a)** Flow cytometry histograms of BV2 cells transfected with pcDNA3 (pcDNA) or pcDNA3-FLAG-PU.1 (FLAG-PU.1) with pCMV-GFP for overexpression and scrambled shRNA (shSCR) or PU.1-targeted shRNA (shA, shB and shD) in pGFP-V-RS vector for knock-down of PU.1 after 3 hours of incubation with red pHrodo-labeled zymosan. Cells were gated on GFP+ populations. **(b)** Flow cytometry analysis of number of gated cells in **a** presented as mean ± SD, pcDNA 67.03 ± 6.883, pcDNA + 1 μM Cyt 15.64 ± 16.24, FLAG-PU.1 82.71 ± 4.74, shSCR 77.17 ± 3.115, shA 48.63 ± 2.285, shB 28.92 ± 2.495, shD 22.76 ± 1.595. pcDNA vs pcDNA + 1 μM Cyt P <0.0001, pcDNA vs FLAG-PU.1 P=0.0306, shSCR vs shA P=0.0002, shSCR vs shB P <0.0001, shSCR vs shD P <0.0001. F(6,13)=58.68, n=3. **(c)** Flow cytometry analysis of geometric mean fluorescent pHrodo intensity in a presented as mean ± SD, pcDNA 10952 ± 2206, pcDNA + 1 μM Cyt 1533 ± 47, FLAG-PU.1 15226 ± 2701, shSCR 13129 ± 4617, shA 9937 ± 2168, shB 8872 ± 2019, shD 8754 ± 1856. pcDNA vs pcDNA + 1 mM Cyt P=0.0092. F(6,13)=6.228, n=3. **(d)** Flow cytometry histograms of BV2 cells transfected as in **(a)** and gated on GFP-populations. **(e)** Flow cytometry analysis of number of gated cells in d presented as mean ± SD, pcDNA 63.92 ± 6.575, pcDNA + 1 μM Cyt 14.21 ± 13.66, FLAG-PU.1 67.54 ± 4.826, shSCR 68.31 ± 5.784, shA 67.27 ± 4.144, shB 65.19 ± 4.268, shD 60.3 ± 2.181. pcDNA vs pcDNA + 1 μM Cyt P <0.0001. F(6,13)=22.53, n=3. **(f)** Flow cytometry analysis of geometric mean fluorescent pHrodo intensity in d presented as mean ± SD, pcDNA 9186 ± 2863, pcDNA + 1 μM Cyt 1545 ± 147, FLAG-PU.1 9931 ± 2458, shSCR 9849 ± 3012, shA 10903 ± 2949, shB 10912 ± 2494, shD 10934 ± 2685. pcDNA vs pcDNA + 1 μM Cyt P=0.0367. F(6,13)=3.473, n=3. **(g)** Phagocytic index of BV2 GFP-cells analyzed in **(e)** and **(f)** presented as mean ± SD, pcDNA 0.5954 ± 0.2223, pcDNA + 1 μM Cyt 0.0209 ± 0.0189, FLAG-PU.1 0.6745 ± 0.188, shSCR 0.6765 ± 0.2274, shA 0.7382 ± 0.2255, shB 0.7131 ± 0.1742, shD 0.6612 ± 0.1748. pcDNA vs pcDNA + 1 μM Cyt P=0.0331. F(6,13)=3.53, n=3. Cytochalasin D treatment in all figures was used as a negative control for phagocytosis. * P <0.05, ** P <0.01, *** P <0.001, repeated measures one-way ANOVA with Sidak’s post hoc multiple comparisons test.

**Supplementary Figure 9.**
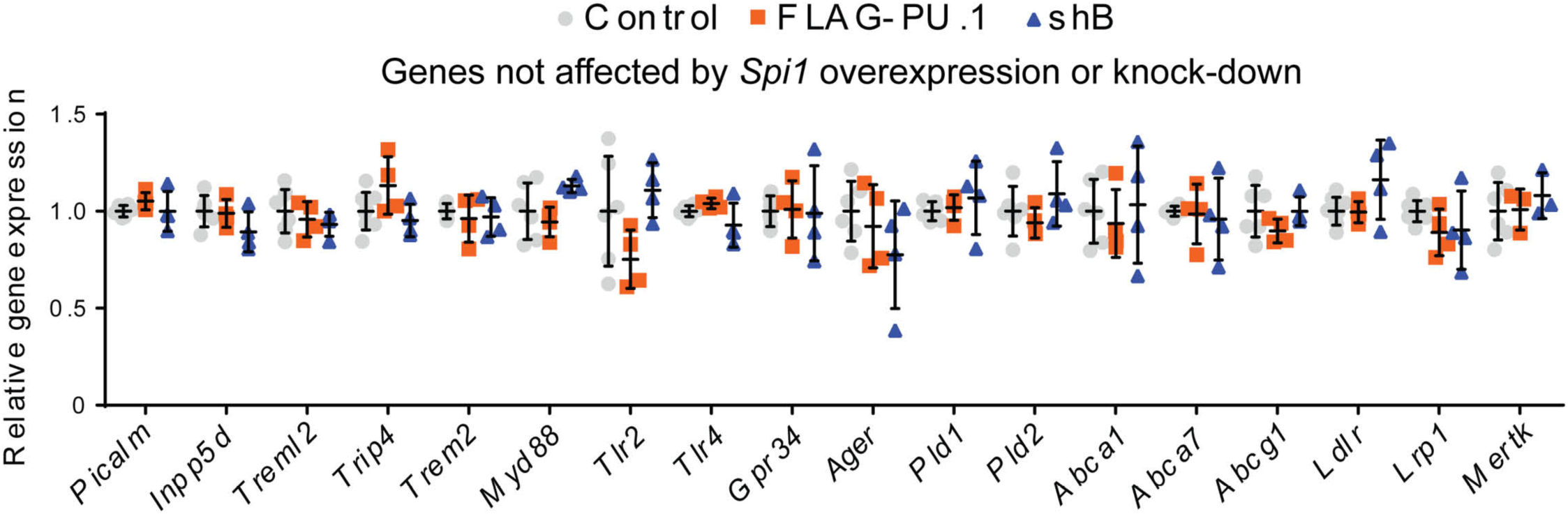
Expression levels of genes related to phagocytosis that were not affected by altered *Spi1* expression. BV2 cells were transiently transfected with pcDNA3-FLAG-PU.1 and pCMV-GFP or pGFP-v-RS-shB against mPU.1. pcDNA3 and pGFP-V-RS-shSCR were used as controls. RNA was extracted from sorted GFP^+^ cells and used for qPCR validation of expression levels for genes of interest. Values are presented as mean ± SD, n=4 samples collected independently.

